# Combined oral vaccination with niche competition can generate sterilizing immunity against entero-pathogenic bacteria

**DOI:** 10.1101/2022.07.20.498444

**Authors:** Verena Lentsch, Aurore Woller, Claudia Moresi, Stefan A. Fattinger, Selma Aslani, Wolf-Dietrich Hardt, Claude Loverdo, Médéric Diard, Emma Slack

## Abstract

Widespread antimicrobial resistance generates an urgent need to develop better disease prophylaxis for intestinal bacterial pathogens. While the first phase of infection with any bacterial pathogen is typically colonization of the mucosal surfaces, current vaccine strategies typically target invasive stages of disease. Here we demonstrate the ability to specifically generate sterilizing immunity against *Salmonella enterica* subspecies *enterica* serovar Typhimurium (*S*.Tm) at the level of gut lumen colonization using a combination of oral vaccination and a rationally-designed niche competitor strain. This is based on the proven ability of specific secretory IgA to generate a fitness disadvantage for a targeted bacterium, allowing a non-targeted competitor to rapidly overtake its niche. By hugely decreasing the population size of an intestinal pathogen during the early stages of infection, this improves protection of gut tissue compared to standard licensed animal vaccines. We demonstrate that most effective protection is generated when the niche competitor is derived from the pathogen and therefore occupies an identical niche. However, as this is unrealistic in real-world infections, we further demonstrate that robust protection can also be generated with a more distantly related “probiotic” niche competitor from a distinct species. Interestingly, focusing prophylaxis on the gut lumen reveals an uncoupling of protective mechanisms required for protection in the gut and gut tissues and those required for protecting against colonization of the spleen and liver. Therefore, while there is still potential to improve this approach by adding systemic immune activation, we nevertheless believe this is a fundamental step forward in our ability to manipulate colonization of intestinal bacteria with potential application to a wide-range of entero-pathogens, as well as to manipulation of microbiota composition.

## Introduction

World-wide, the frequency of drug-resistant infections with *E. coli* and with non-typhoidal *Salmonella* have been steadily increasing, posing a looming challenge for health systems^1^. There is a pressing need to develop control and prevention strategies that are independent of antibiotics. Vaccination of both farm animals and humans is a promising alternative. While high-efficacy vaccine candidates against Typhoid fever are currently well advanced (e.g. intramuscular Vi capsule, typhoid conjugate vaccines Vi-TT, and oral Ty21a live-attenuated vaccines)^2,3^, there are no licensed human vaccines for non-typhoidal Salmonellosis (NTS)^4^. Moreover, licensed live-attenuated *S*.Tm swine and bovine vaccines show only weak protection from colonization^5^ and can cause disease in highly-susceptible individuals, such that versions currently licensed for animal use are not directly suitable for human translation^6^. An ideal solution would be fully inactivated vaccines that can mimic the immunity and protection generated by live-attenuated vaccines but with a much higher safety profile and robustness.

When designing vaccines targeting intestinal bacteria, we have either focused on mimicking natural infection^7^ or on generating immune responses targeting specific pathogen antigens^8,9^. However, we increasingly realize that infection of the gut involves disruption of a densely populated microbial ecosystem, and the intestinal microbiome is itself part of our defenses^10^. Therefore, evidence suggests that naturally arising immunity at our mucosal surfaces does not operate in isolation, but rather interacts with the endogenous microbiota to recover homeostasis and evict colonizing opportunistic pathogens^11^. For example, fecal microbiota transplantation can be curative in recurrent *Clostridioides difficile* infections^12^, and introduction of the pneumococcal conjugate vaccines has resulted in displacement of vaccine-targeted *Streptococcus pneumoniae* serovars from the nasopharynx by non-targeted strains in both vaccinated as well as unvaccinated individuals^13,14^.

To prevent tissue invasion in hosts permissive for systemic *S*.Tm growth, inflammation and systemic spread, we have previously estimated that a >1000-fold reduction in invasion rate of *S*.Tm is required during the first day of infection^15^. The overall invasion rate is a product of the gut luminal *S*.Tm population size and the ability of each individual bacterium to invade. Antibody-mediated clumping, via enchained growth and/or agglutination achieves around a 100-fold reduction in the planktonic infectious population size^16^, but alone cannot be sufficient to fully prevent invasive disease. Therefore, vaccination strategies are required that more efficiently prevent colonization of the gut lumen.

At the simplest level, the population size of *S*.Tm in the gut lumen is determined by its growth rate, clearance rate and the size of the available metabolic niche. The available metabolic niche can be dramatically shrunk by competitors using the same resources. A classic example of this phenomenon occurs when “cheater” mutations spontaneously arise during chronic NTS^17^. Mutations in the *Salmonella* Pathogenicity Island 1 (SPI-1) master-regulator of virulence *hilD* can outgrow and displace wildtype Salmonella from the gut lumen. This is attributed to a fast net-growth rate and increased stress resistance of *hilD*-mutant strains due to loss of the fitness costs associated with expression of the HilD regulon^18,19^.

Niche competition can also be favoured by increasing the clearance rate of the pathogen. High-affinity intestinal IgA, induced by oral vaccination with whole-cell inactivated vaccines, can increase *S*.Tm clearance rates by generating bacterial clumps (via enchained growth or agglutination) that are more efficiently cleared in the flow of the fecal stream^16^. We have recently demonstrated that this IgA-mediated selective pressure can be used to manipulate the evolutionary trajectory of *Salmonella enterica* subspecies *enterica* serovar Typhimurium (*S*.Tm) in the gut lumen^20^. Moreover, part of this work demonstrated the ability of IgA to generate a more than a million-fold ratio of a targeted *S*.Tm strain over a non-bound strain^20^. Therefore, IgA protects the intestinal environment not only via immune exclusion (i.e. preventing interaction of pathogenic bacteria with the epithelium) but also by competitively eliminating targeted bacteria from the intestinal ecosystem. This suggested a huge potential for specific high-affinity IgA responses to manipulate the outcome of competition between an invading pathogen and an engineered niche competitor.

Critical components of such a prophylactic system are: 1) a vaccine capable of inducing high-affinity specific IgA against the pathogen of interest; 2) a non-pathogenic strain with (ideally) complete metabolic niche overlap with the pathogen of interest, a faster growth-rate than the pathogen of interest and absence of surface antigen cross-reactivity to the pathogen of interest. Additionally, questions arise as to how vaccines and competitors can be combined temporally to give maximum effect, and the extent of protection from invasive disease that can be achieved with such an approach. Here we make use of the well-established and severe, murine model of non-typhoidal Salmonellosis to establish this concept. In this model, oral antibiotics are applied to acutely generate a large open niche for *S*.Tm in the mouse cecum and upper large intestine^21^. Subsequently, very low numbers of *S*.Tm can be orally inoculated and will rapidly grow to fill the available niche. Disease depends on the activity of *Salmonella* Pathogenicity Islands 1 and 2 and includes acute typhlocolitis, colonization of the intestinal tissue, mesenteric lymph nodes, spleen and liver^21-23^. In the resistant (Nramp1+/+, also known as Slc11a1) 129SJL mouse strain, the disease is slowly controlled, but full recovery takes more than 1 month^24^. We also extend our observations to the murine oral Typhoid fever model, in which a high-dose of *S*.Tm is delivered orally to mice with an intact intestinal microbiota, resulting in a disease that is more reliant on the tissue-invasive stages of disease.

Here we demonstrate that combining a benign niche-competitor and a whole-cell inactivated vaccination regimen can profoundly suppress intestinal colonization with virulent *Salmonella* even on high-dose exposure in a very high-susceptibility model. Protection from gut tissue invasion and inflammation was also robust. Interestingly, live-attenuated *Salmonella* vaccines were superior to inactivated vaccines in protecting from colonization of systemic sites, indicating that mechanisms that protect systemic sites are discrete from intestinal protection.

## Results

### Combining oral inactivated bacterial vaccination with a bacterial niche competitor protects mice from intestinal inflammation and leads to rapid *S*.Tm^WT^ clearance in a model of NTS

As a first proof-of-principle we made use of *S*.Tm carrying a mutation in the “Salmonella Pathogenicity Island 1” (SPI-1) master regulator *hilD*, that has previously been demonstrated to outgrow wild-type *S*.Tm in the gut lumen during long-term infections^18,25^. This was combined with inactivating mutations in *ssaV* to disrupt the function of “Salmonella Pathogenicity Island 2” (SPI-2) and deletion of *oafA* to prevent acetylation of the *S*.Tm O-antigen, i.e. preventing generation of the O:5 epitope. The resulting strain is fully avirulent^18^ (**Fig. S1**), fast-growing^18^, and less bound by IgA induced by a wild-type O:5,12-0 *Salmonella* vaccine than the isogenic wild-type strain^20^. The resulting mutant *S*.Tm^*hilD ssaV oafA*^ was used as niche competitor hereafter named *S*.Tm^Comp^.

To demonstrate the effect of pathogen-targeting IgA on competition, SPF 129S6/SvEv wildtype mice received an inactivated whole-cell oral *S*.Tm vaccine once weekly for 4 weeks or a mock PBS treatment. Subsequently, mice were antibiotic treated to eliminate a large part of the microbiota and infected with either virulent wildtype *S*.Tm (*S*.Tm^WT^) alone, or *S*.Tm^WT^ combined 1:1 with *S*.Tm^Comp^. Intestinal colonization and inflammation were monitored for 10 days post infection and tissue invasion and histopathology were monitored at endpoint (**Fig. 1A**).

**Figure 1.**
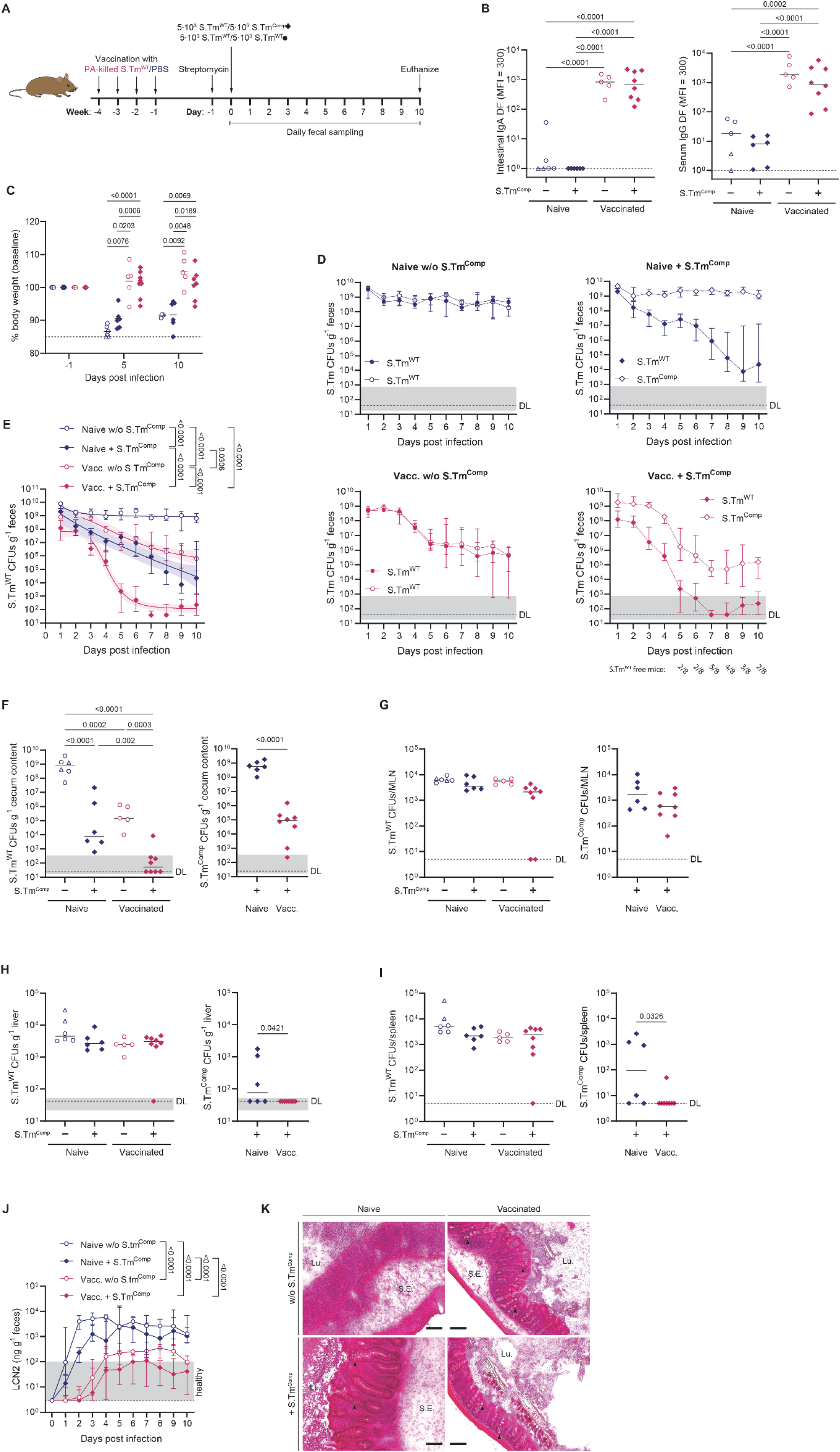
Combining oral vaccination with niche competition protects mice from intestinal inflammation and leads to rapid *S*.Tm^WT^ clearance. PBS (blue symbols) or PA-*S*.Tm-vaccinated (pink symbols) 129S6/SvEv mice were pretreated with streptomycin and infected with a total of 10^4^ of a 1:1 mixture of two isogenic *S*.Tm^WT^ strains (open circles) or *S*.Tm^WT^ and *S*.Tm^Comp^ (*S*.Tm^*hilD ssaV oafA*^, filled diamonds). (**A**) Experimental procedure. (**B**) *S*.Tm^WT^ (O:4[5]12-0) specific intestinal IgA and serum IgG titres as determined by flow cytometry. (**C**) Change in body weight over the course of infection. Solid lines show the mean. (**D**) Fecal CFUs as determined by selective plating. Open and filled symbols represent either isogenic *S*.Tm^WT^ strains (circles) or *S*.Tm^WT^ and *S*.Tm^Comp^ (diamonds) with different antibiotic resistances. (**E**) A 4-parameter logistic curve fit was fitted to the fecal *S*.Tm^WT^ CFUs and the area under the curve was statistically compared. Shaded areas depict the 95% CI of the fits. (**F-I**) *S*.Tm CFUs in cecum content (**F**), MLN (**G**), liver (**H**) and spleen (**I**). (**J**) Intestinal inflammation was determined by measuring fecal lipocalin-2. (**K**) Histology of cryo-embedded H&E-stained cecal tissue sections. Arrowheads show exemplary goblet cells. Scale bars: 100 µm. Pooled data from two independent experiments with switched antibiotic resistances (n = 5-8 mice/group). Solid lines depict the median unless stated otherwise, error bars the interquartile range. Dotted lines show the detection limit and the shaded area the range for cases in which the detection limit is dependent on sample weight. Open triangles show mice that had to be euthanized prematurely due to excessive weight loss (≥ 15%) or disease symptoms. Statistics were performed by mixed-effects analysis (C) or one-way analysis of variance (ANOVA) (C) on log-normalized data (B, F-I) or area under the curve (AUC) (E, J). Where only two groups were compared, an unpaired two-tailed t-test on log-normalized data was done (F-I). CFU, colony forming unit; DF, dilution factor; LCN2, lipocalin-2; Lu., Lumen; MFI, median fluorescence intensity; MLN, mesenteric lymph node, S.E., submucosal edema.

In line with published data, we could detect high levels of *S*.Tm O:5,12-0-specific IgA in the intestine as well as IgG in serum of vaccinated mice at endpoint, regardless of the subsequent infection (**Fig. 1B**). This *S*.Tm specific antibody response was able to protect the vaccinated mice from weight loss upon infection with WT *S*.Tm (**Fig. 1C**). In contrast, all mice that were not vaccinated experienced significant weight loss until 5 days post infection. One unvaccinated mouse of each of two independent experiments had to be euthanized at 5 days post infection due to excessive weight loss.

As antibiotic pre-treatment in this model generates a huge empty niche for *S*.Tm colonization in the gut lumen, *in vivo* binding of IgA to *S*.Tm^WT^ could not prevent initial expansion of *S*.Tm, and total *S*.Tm counts were similar in all groups. However, over time the differences between groups become clear: *S*.Tm^WT^ CFU/g feces remains constant over 10 days in unvaccinated mice without *S*.Tm^Comp^. The presence of either vaccination alone, or *S*.Tm^Comp^ results in a drop in *S*.Tm^WT^ CFU from day 2-3 post infection, with CFU 10^5^ CFU/g feces at day 10. In contrast, the combination of vaccination and *S*.Tm^Comp^ generates an exponential drop in fecal CFU of *S*.Tm^WT^ over the first 5 days of infection, with some mice completely clearing the pathogenic strains by day 5 (**Fig. 1D and E**). The cecum content counts reflected the picture found in feces with a much lower burden of *S*.Tm^WT^ in all treated groups and absence of *S*.Tm^WT^ in 50% of the mice that received vaccination and the niche competitor (**Fig. 1F**). Interestingly, despite a very dramatic clearance of *S*.Tm^WT^ from the gut lumen, colonization of systemic sites was apparently unaltered by either vaccination or by introduction of the niche competitor (**Fig. 1G-I**). This is consistent with quantitative modelling of tissue invasion in murine NTS, which predicted that a more than 1000-fold reduction in *Salmonella* intestinal load is required to completely inhibit systemic spread^15,16^. As it takes several days for the *S*.Tm^WT^ levels to be reduced below this level in our treatment groups, there remains a considerable time-window for systemic spread to occur. While we cannot exclude that differences may have been more apparent at earlier time-points, this clearly indicates that encounter of a niche competitor simultaneous to infection, even in orally vaccinated mice that display both intestinal IgG and serum IgA responses, is not sufficient to prevent colonization of the spleen and liver of mice.

The partial-protective capacity of IgA with and without niche competition was also apparent when looking at intestinal inflammation as measured by lipocalin-2 levels in feces. In both vaccinated groups, inflammation not only started later but also the maximum fecal lipocalin-2 was significantly lower throughout the whole course of infection (**Fig. 1J**). This was underpinned by histological stainings at day 10 post challenge that showed no pathological changes in mice that were both vaccinated and colonized with *S*.Tm^Comp^, and only mild changes in the vaccine-alone group (**Fig. 1K**). Intestinal inflammation could not be significantly prevented by the presence of the niche-competitor alone when it is introduced simultaneously with the infectious challenge (**Fig. 1J and K**).

These experiments indicated the feasibility of combining vaccination and niche competition, while revealing the need to optimize the procedure for protection of both the gut and systemic sites.

### Modelling the interaction of niche competition and vaccination

As intestinal colonization is necessarily a highly dynamic process, we built a simple mathematical model to generate predictions on the requirements for extinction of *S*.Tm^WT^ and the time-to-extinction. We predict that minimizing the time-to-extinction in the gut lumen will best inhibit systemic spread of *S*.Tm and can minimize the risk of immune escape.

In this model we assumed that the microbiota (of size M(t)), *S*.Tm^Comp^ (of size C(t)) and *S*.Tm^WT^ (of size W(t)) compete for undefined shared nutrient resources, which leads to a carrying capacity K_1_. Additionally, we assumed a second independent nutrient source for *S*.Tm^Comp^ and *S*.Tm^WT^ on the one hand, and for the microbiota on the other hand leading to carrying capacities K_2_ and K_3_. In a deterministic view, the size of the populations evolves as

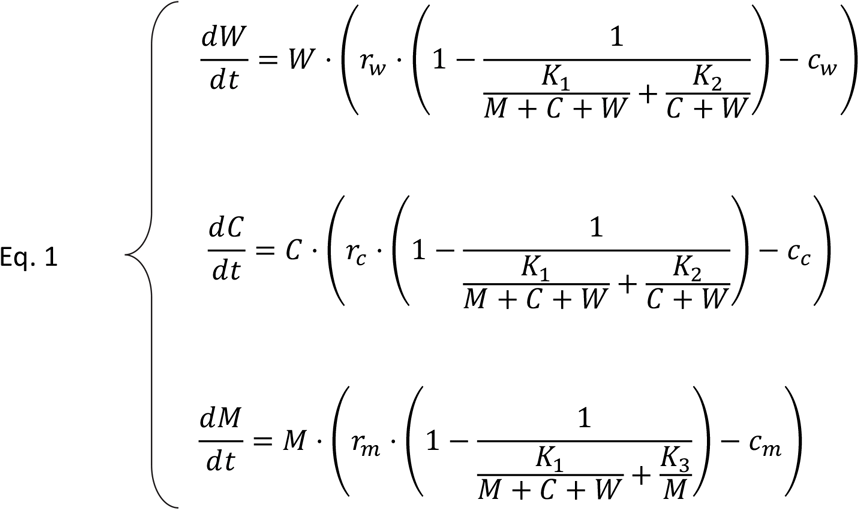

where r_w_, r_c_ and r_m_ correspond to the growth rate of *S*.Tm^WT^, *S*.Tm^Comp^ and the microbiota, respectively. The parameters c_w_, c_c_ and c_m_ correspond to the clearance rates of these populations.

In order to predict “extinction probability” and “extinction time”, we required realistic estimates of the population dynamics of the two *S*.Tm strains and the “average microbiome”. *S*.Tm population dynamics parameters in the presence or absence of IgA were estimated by fitting the competition data shown in Figure 1 and Figure S2A (see materials and methods) (**Fig. 2A**). This confirmed the predicted higher net growth rate for *S*.Tm^Comp^ as compared to *S*.Tm^WT^ and the elevated clearance rate of *S*.Tm^WT^ in vaccinated mice (see **Table 1**).

**Figure 2:**
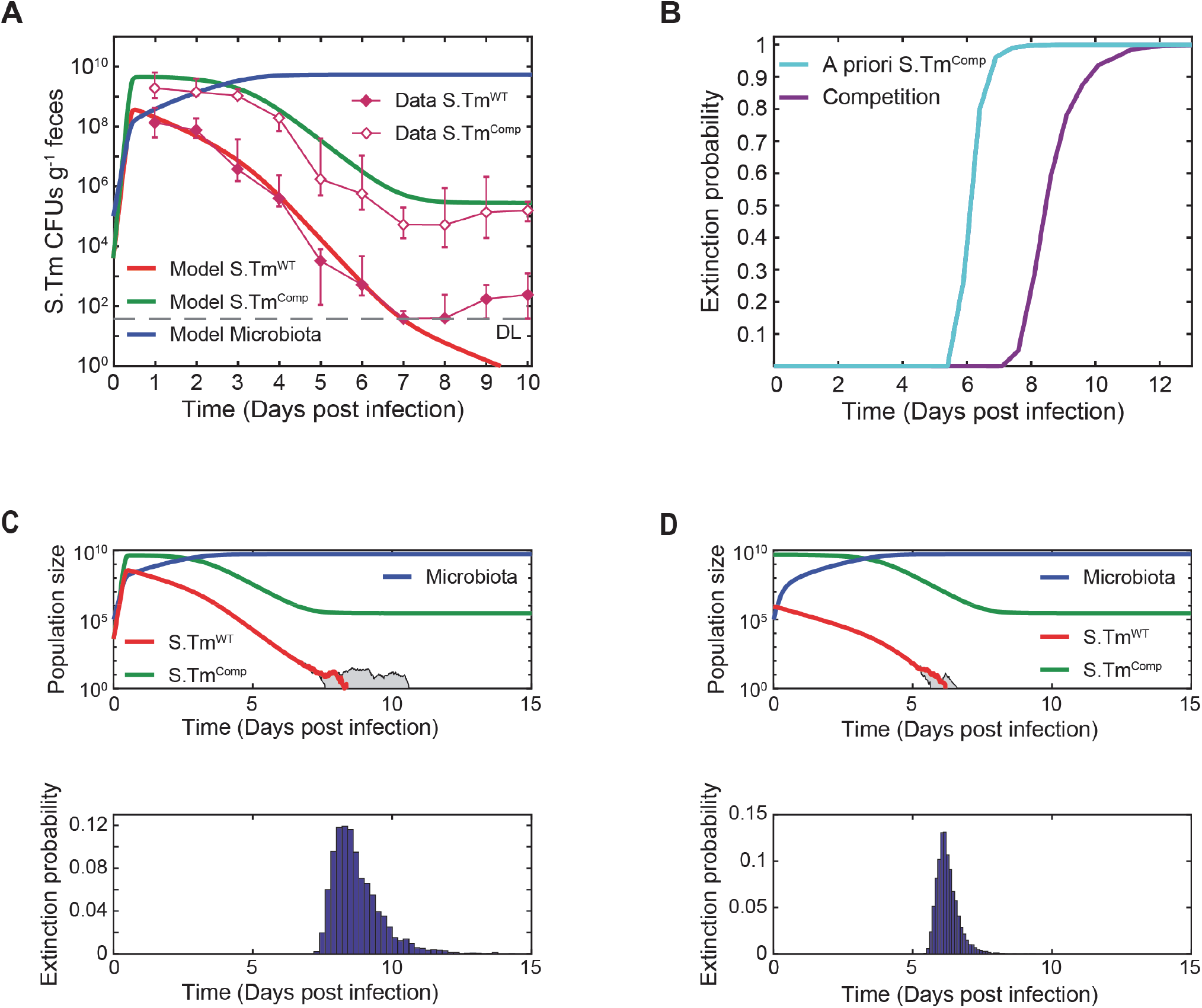
Modelling of *S*.Tm^WT^ extinction for vaccination and/or niche competition. (**A**) Adjustment of the model to the data shown in Figure 1. The thick lines correspond to the prediction from the model and the thin lines with diamonds to the experimental data (median ± interquartile range). Initial *S*.Tm^WT^ and *S*.Tm^Comp^ CFUs: 4·10^3^. (**B**) Extinction probability of *S*.Tm^WT^ over time for vaccinated mice + *S*.Tm^Comp^. The purple line shows the prediction of the model when *S*.Tm^WT^ and *S*.Tm^Comp^ are given at the same time, the turqoise line for the case that *S*.Tm^Comp^ is given 3 days prior to *S*.Tm^WT^. The simulations were performed with the deterministic model for large population size and with the Gillespie algorithm (1000 realizations) for small size of *S*.Tm^WT^ population (*S*.Tm^WT^ < 100). (**C**) Extinction time probability distribution for *S*.Tm^WT^ and *S*.Tm^Comp^ given at the same time. Simulations were performed with the deterministic model for large population size. For small size of *S*.Tm^WT^ population (*S*.Tm^WT^ < 100), simulations were performed with the Gillespie algorithm (5000 realizations). Upper panel: the red line corresponds to a trajectory whose extinction time is around the median extinction time. The gray area shows trajectories of the 2.5^th^ percentile extinction time and the 97.5^th^ percentile extinction time. Lower panel: extinction time probability distribution. The y-axis corresponds to (bin height*bin width). Median extinction time: 8.5 days. 2.5^th^ percentile: 7.6 days. 97.5^th^ percentile: 11 days. Initial *S*.Tm^WT^ CFUs: 4·10^3^. Initial *S*.Tm^Comp^ CFUs: 4·10^3^. (**D**) Extinction time probability distribution for a priori colonization with *S*.Tm^Comp^. Simulations were performed with the deterministic model for large population size. For small size of *S*.Tm^WT^ population (*S*.Tm^WT^ < 100), simulations were performed with the Gillespie algorithm (5000 realizations). Upper panel: the red line corresponds to a trajectory whose extinction time is around the median extinction time. The gray area shows trajectories of the 2.5^th^ percentile extinction time and the 97.5^th^ percentile extinction time. Lower panel: extinction time probability distribution. The y-axis corresponds to (bin height*bin width). Median extinction time: 6 days. 2.5^th^ percentile: 5.7 days. 97.5^th^ percentile: 7.2 days. Initial *S*.Tm^WT^ CFUs: 5·10^9^. Initial *S*.Tm^Comp^ CFUs: 4·10^3^. The dotted line depicts the mean detection limit of the experimental data. (**A, C, D**) Parameter values: r_w_ = 33.7 day^-1^, c_w_ = 6.4 day^-1^, r_c_ = 38.7 day^-1^, c_c_ = 5.9 day^-1^, r_m_ = 18.7 day^-1^, c_m_ = 1.7 day^-1^, K_1_ = 5.9·10^9^, K_2_ = 2.3·10^4^, K_3_ = 10^7^ and m_0_ = 10^5^. r_w_, r_c_ and r_m_: growth rate of *S*.Tm^WT^, *S*.Tm^Comp^ and the microbiota, respectively. c_w_, c_c_ and c_m_: clearance rates of *S*.Tm^WT^, *S*.Tm^Comp^ and the microbiota. K_1_, K_2_, K_3_: carrying capacities of *S*.Tm and the microbiota, *S*.Tm^WT^ and *S*.Tm^Comp^ and the microbiota alone. m_0_: size of microbiota after antibiotic clearance.

**Table 1:** Kinetic parameter values inferred from the competition data shown in Fig. 1 (for vaccinated + *S*.Tm^Comp^ group). r_w_, r_c_ and r_m_: growth rate of *S*.Tm^WT^, *S*.Tm^Comp^ and the microbiota, respectively. c_w_, c_c_ and c_m_: clearance rates of *S*.Tm^WT^, *S*.Tm^Comp^ and the microbiota. K_1_, K_2_, K_3_: carrying capacities of *S*.Tm and the microbiota, *S*.Tm^WT^ and *S*.Tm^Comp^ and the microbiota alone. m_0_: size of microbiota after antibiotic clearance at time point of infection with *S*.Tm^WT^.

| $r_w$ | $r_c$ | $r_m$ | |
| --- | --- | --- | --- |
| 33.7 day<sup>-1</sup> | 38.7 day<sup>-1</sup> | 18.7 day<sup>-1</sup> |  |
| $c_w$ | $c_c$ | $c_m$ | |
| 6.4 day<sup>-1</sup> | 5.9 day<sup>-1</sup> | 1.7 day<sup>-1</sup> |  |
| $K_1$ | $K_2$ | $K_3$ | $m_0$ |
| $5.9 \cdot 10^9$ | $2.3 \cdot 10^4$ | $10^7$ | $10^5$ |

Using 2500 iterations of a stochastic version of this model, we could derive extinction-related parameters. Based on continuous growth and clearance of the gut luminal *Salmonella* population, these models indicate clearance of virulent *Salmonella* within eight days of infection in the mice treated with vaccination and *S*.Tm^Comp^, compared to continuous colonization in mice untreated or receiving vaccine only (**Fig. 2B and C**). Interestingly, the model predicted that the time-point for introduction of the niche competitor was not a major variable, as long as this occurred prior to infection. Major determinants of success are the relative growth rates and loss rates of the competitor versus the virulent bacterium as well as the carrying capacity, which is in turn determined by the regrowth of the microbiota. Therefore, a major conclusion of the model is that the niche competitor should be already present in the gut for effective protection (**Fig. 2B and D**), which is also a more realistic situation for disease prophylaxis.

Longer duration of infection and more intense *Salmonella* shedding increase the probability of transmission^26^. Both can be dramatically reduced by vaccination/competition, and the model reveals a major potential for this intervention to reduce the *Salmonella* burden in farm animals.

### Introduction of the niche competitor a priori allows complete pathogen clearance when combined with oral vaccination

Based on the modelling data, we carried out a vaccination and challenge experiment, this time introducing the competitor strain carrying the same Streptomycin resistance as *S*.Tm^WT^ three days prior to infection (**Fig. 3A**). We additionally increased our *S*.Tm^WT^ infection dose to 1·10^6^ CFU, in order to increase the stringency of the challenge. Again, we could confirm that vaccination leads to a robust induction of *S*.Tm-specific intestinal IgA as well as serum IgG (**Fig. 3B**), and this protected animals from weight loss upon *S*.Tm^WT^ infection, regardless of the higher dose. In contrast to simultaneous infection, introducing *S*.Tm^Comp^ a priori into naïve mice also prevented weight loss (**Fig. 3C**). In animals infected with *S*.Tm^WT^ only, we observe very similar infection kinetics to those induced by challenging with 5·10^3^ CFU. By contrast, *S*.Tm^WT^ struggled to expand at all in mice already colonized with the competitor(**Fig. 3D**) and was eliminated as soon as four days post infection in mice protected by both vaccination and pre-colonization with *S*.Tm^Comp^. *S*.Tm^WT^ was undetectable in the cecum content in 5 of 8 “vaccinated + *S*.Tm^Comp^ colonized” mice on the final day of infection (**Fig. 3E**). This fitted well to predictions from the model derived above (**Fig. 3F**). The predictions of the model were less accurate for *S*.Tm^WT^ at later time points, which might be due to the emergence of escape mutants. Intestinal inflammation could also be completely prevented by the combined prophylaxis, remaining within the range of healthy animals for the full duration of the infection (**Fig. 3G-I**). Moreover, systemic spread of *S*.Tm^WT^ could be effectively prevented by the combination of vaccination and *S*.Tm^Comp^ in around 50% of treated animals (**Fig. 3J-L**), and correlated well with early loss of *S*.Tm^WT^ from the gut, rendering some mice completely pathogen free at 10 days post infection. Therefore, combining an inactivated oral vaccine with a live niche competitor can generate sterilizing immunity against virulent *Salmonella* in a very severe model of non-typhoidal Salmonellosis.

**Figure 3.**
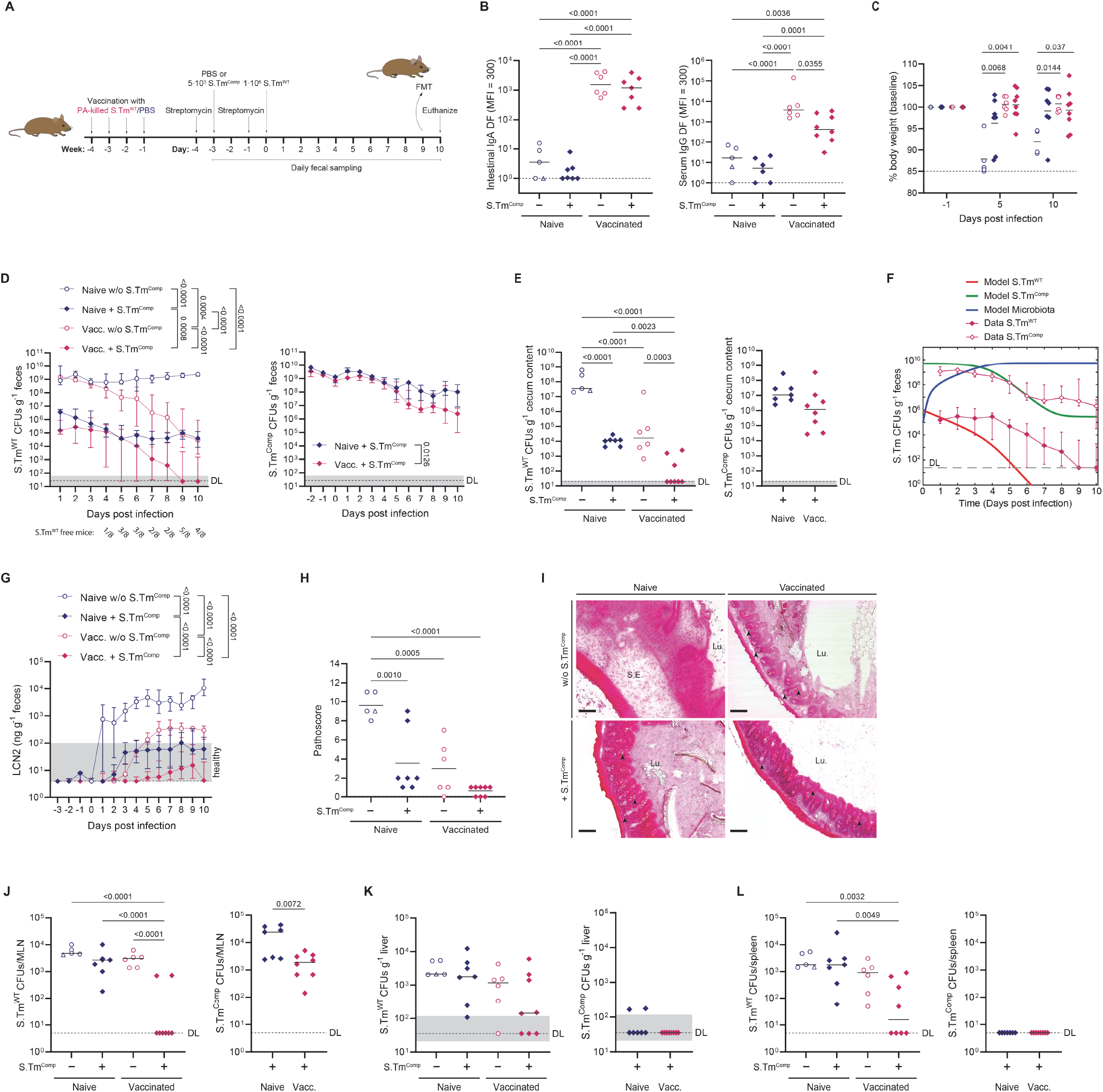
A priori colonization with *S*.Tm^Comp^ can generate sterilizing immunity in the gut when combined with vaccination. PBS (blue symbols) or PA-*S*.Tm-vaccinated (pink symbols) 129S6/SvEv mice were pretreated with streptomycin and infected with 1·10^6^ *S*.Tm^WT^. Two groups were pre-colonized with 5·10^3^ *S*.Tm^Comp^ 3 days before infection. (**A**) Experimental procedure. (**B**) *S*.Tm^WT^-specific intestinal IgA and serum IgG titres as determined by flow cytometry. (**C**) Change in body weight over the course of infection. Solid lines depict the mean. Fecal (**D**) and cecum content (**E**) *S*.Tm CFUs as determined by selective plating. (**F**) Comparison of the predictions of the model to the experimental data. The thick solid lines correspond to the prediction from the model, using the value of the dynamical parameter inferred from the experiment with simultaneous *S*.Tm^WT^ and *S*.Tm^Comp^ infection. The thin lines with diamonds to the experimental data (median ± interquartile range). Initial *S*.Tm^WT^ CFUs: 5·10^9^. Initial *S*.Tm^Comp^ CFUs: 4·10^3^. Parameter values: r_w_ = 33.7 day^-1^, c_w_ = 6.4 day^-1^, r_c_ = 38.7 day^-1^, c_c_ = 5·9 day^-1^, r_m_ = 18.7 day^-1^, c_m_ = 1.7 day^-1^, K_1_ = 5.9.10^9^, K_2_ = 2.3·10^4^, K_3_ = 10^7^ and m_0_ = 10^5^. r_w_, r_c_ and r_m_: growth rate of *S*.Tm^WT^, *S*.Tm^Comp^ and the microbiota, respectively. c_w_, c_c_ and c_m_: clearance rates of *S*.Tm^WT^, *S*.Tm^Comp^ and the microbiota. K_1_, K_2_, K_3_: carrying capacities of *S*.Tm and the microbiota, *S*.Tm^WT^ and *S*.Tm^Comp^ and the microbiota alone. m_0_: size of microbiota after antibiotic clearance. (**G-I**) Intestinal inflammation as determined by fecal lipocalin-2 (**G**) and histopathological scoring of cecal tissue sections (**H**) Solid lines in (H) show the mean. (**I**) Representative images of H&E-stained cecal tissue sections. Arrowheads show exemplary goblet cells. Scale bars: 100 µm. (**J-L**) *S*.Tm CFUs in MLN (**J**), liver (**K**) and spleen (**L**). Pooled data from two independent experiments with switched antibiotic resistances (n = 5-8 mice/group). Solid lines depict the median unless stated otherwise, error bars the interquartile range. Dotted lines show the detection limit and the shaded area the range for cases in which the detection limit is dependent on sample weight. Open triangles show mice that had to be euthanized prematurely due to excessive weight loss (≥ 15%) or disease symptoms. Statistics were performed by mixed-effects analysis (C) or one-way ANOVA (C, G) on log-normalized data (B, E, I-K) or area under the curve (AUC) (D, F). Where only two groups were compared, an unpaired two-tailed t-test on log-normalized data was done (E, I-K). CFU, colony forming unit; DF, dilution factor; FMT, fecal microbial transplant; LCN2, lipocalin-2; Lu., lumen; MFI, median fluorescence intensity; MLN, mesenteric lymph node; S.E., submucosal edema.

To further corroborate our findings of sterilizing immunity, we performed fecal microbial transplants from vaccinated/*S*.Tm^Comp^ mice into naive Streptomycin pre-treated 129S6/SvEv mice. All but one mouse from this group, did not efficiently transmit *S*.Tm^WT^ by fecal microbial transplantation (FMT) (**Fig. S3**). In contrast, all untreated control mice used for this experiment efficiently transferred *S*.Tm^WT^ and disease to the naïve mice (**Fig. S3**).

In summary, we could show that our niche competitor when given a priori is able to establish a partial colonization resistance thereby limiting intestinal *S*.Tm^WT^ expansion. When combined with vaccination, a major fraction of the animals could completely clear *S*.Tm^WT^ from all examined sites i.e. generating sterilizing immunity. Importantly, we could not only greatly ameliorate disease in animals treated with this method, but we could also prevent transmission in almost 90% of the cases.

### Commensal competitors and vaccination

While the combination of inactivated oral vaccines and an engineered *Salmonella*-derived competitor is highly effective, there remain potential safety/legislative issues associated with the application of live GMOs and a very low, but non-zero, risk of reversion of the competitor strain via horizontal gene transfer^27^. An attractive alternative would be to develop a non-pathogenic commensal bacterial strain that can nevertheless compete efficiently with *S*.Tm^WT^. For this purpose, we chose the mouse commensal ECOR B2 strain *E. coli* 8178 (Ec^8178^). This has been demonstrated to grow well in *S*.Tm-permissive conditions^28^ and could limit *S*.Tm infection after dietary perturbation^29^. Moreover, this *E. coli* produces an unrelated O-antigen structure, allowing us to use an “evolutionary trap” version^20^ of our *S*.Tm vaccine covering all possible variations of the *S*.Tm O-antigen. This adds robustness to the technique as selective pressure exerted by vaccine-induced IgA can provide a selective advantage both to the competitor strain, and to naturally emerging *S*.Tm variants with a short O-antigen. As mutants with short LPS are susceptible to complement, bile acids and other membrane-targeting stresses, this is associated with decreased virulence of *S*.Tm^WT^ in vaccinated mice^20^.

We therefore repeated our previous experiment using the evolutionary trap (EvoTrap) vaccine together with Ec^8178^ as a competitor given three days prior to challenge (**Fig. 4A**). Vaccination with the EvoTrap vaccine led to high levels of *S*.Tm O:5,12-0 specific antibodies in small intestine and blood (**Fig. 4B**) as well as to the other *S*.Tm O-antigen variants (**Fig. S4**). Similarly, to what we have observed with giving *S*.Tm^Comp^ prior to *S*.Tm^WT^ infection, also Ec^8178^ alone could reduce weight loss after challenge and weight loss was prevented in both vaccinated groups (**Fig. 4C**). Ec^8178^ colonized the mouse gut to high levels and was able to decrease initial *S*.Tm^WT^ expansion. However, Ec^8178^ was clearly less effective as a competitor in the intestine when compared with *S*.Tm^Comp^, as *S*.Tm^WT^ levels remained high (>10^6^) in feces and cecum of all groups at day 10 post infection (**Fig. 4D and E**). Despite the high *S*.Tm^WT^ numbers in the gut, we observed a significant reduction in systemic counts (mesenteric lymph nodes, spleen and liver) in the vaccinated group that received Ec^8178^ (**Fig. 4F-H**). When analysing intestinal inflammation, we again found that the intestinal inflammation marker lipocalin-2 was almost completely absent from the “vaccinated + Ec^8178^ pre-colonized” group, indicating robust protection from tissue invasion and disease, despite incomplete clearance of *S*.Tm^WT^ from the gut lumen (**Fig. 4I-K**). As predicted^20^ with EvoTrap vaccination, a major fraction of luminal *S*.Tm^WT^ re-isolated from vaccinated mice carried a spontaneous deletion resulting in loss of *wzyB* and therefore short O-antigen production, which may explain this discrepancy (**Fig. S5**).

**Figure 4.**
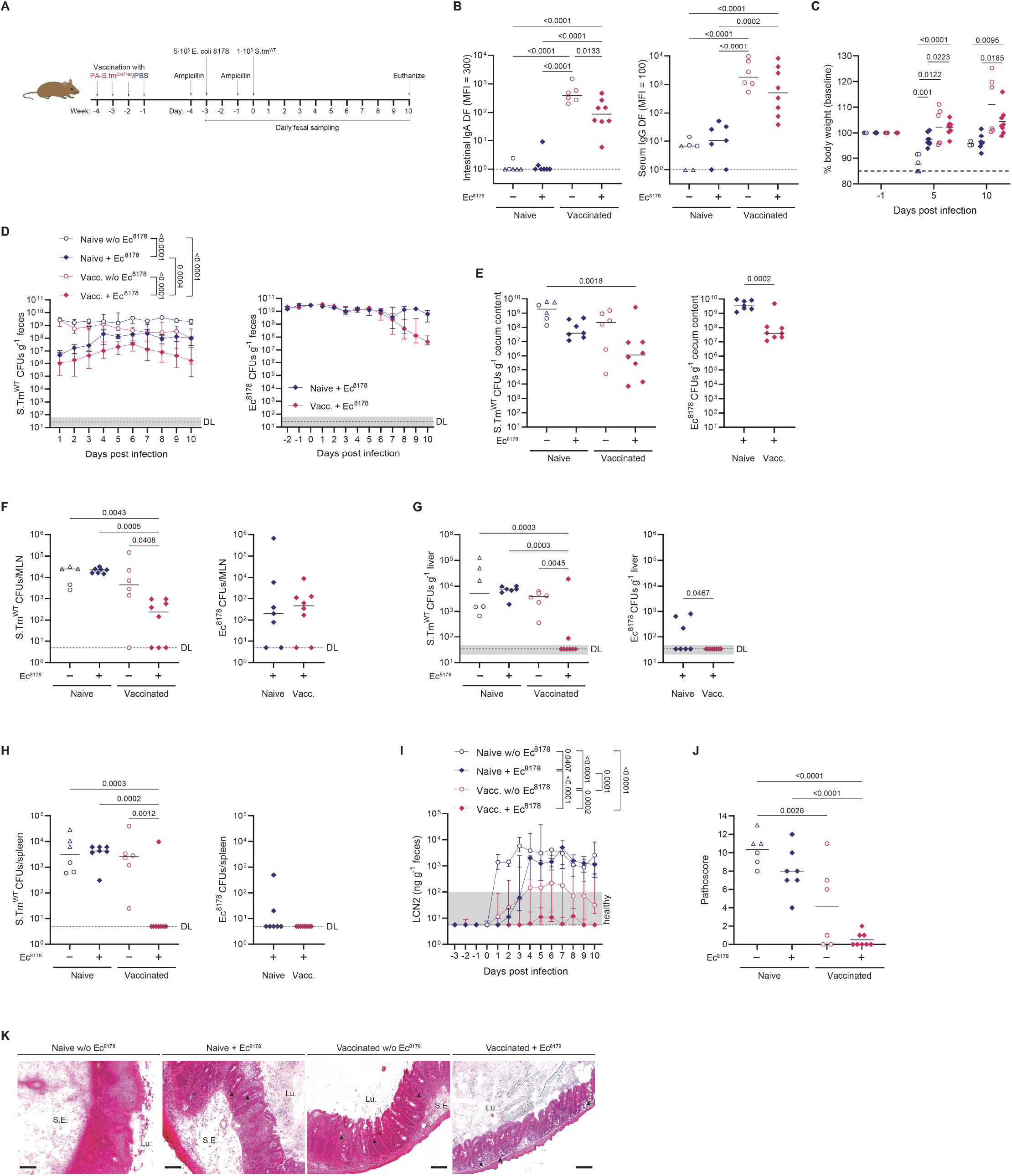
Vaccination together with the commensal competitor *E. coli* 8178 can prevent intestinal pathology albeit *S*.Tm^WT^ cannot be cleared from the gut. PBS (blue symbols) or EvoTrap-vaccinated (pink symbols) 129S6/SvEv mice were pretreated with ampicillin and infected with 1·10^6^ *S*.Tm^WT^. Two groups were pre-colonized with 5·10^3^ *E. coli* 8178 3 days before infection. (**A**) Experimental procedure. (**B**) *S*.Tm^WT^-specific intestinal IgA and serum IgG titres as determined by flow cytometry. (**C**) Change in body weight over the course of infection. Solid lines depict the mean. (**D-H**) CFUs in feces (**D**) cecum content (**E**), MLN (**F**), liver (**G**) and spleen (**H)**. Intestinal inflammation was determined by fecal lipocalin-2 (**I**) and histopathological scoring of cecal tissue sections (**J**). Solid lines in (J) show the mean. (**K**) Representative images of H&E-stained cecal tissue sections. Arrowheads show exemplary goblet cells. Scale bars: 100 µm. Pooled data from two independent experiments (n = 6-8 mice/group). Solid lines depict the median unless stated otherwise, error bars the interquartile range. Dotted lines show the detection limit and the shaded area the range for cases in which the detection limit is dependent on sample weight. Open triangles show mice that had to be euthanized prematurely due to excessive weight loss (≥ 15%) or disease symptoms. Statistics were performed by mixed-effects analysis (C) or one-way ANOVA (C, J) on log-normalized data (B, E-H) or area under the curve (AUC) (D, I). Where only two groups were compared, an unpaired two-tailed t-test on log-normalized data was done (E-H). CFU, colony forming unit; DF, dilution factor; LCN2, lipocalin-2; Lu., lumen; MFI, median fluorescence intensity; MLN, mesenteric lymph node; S.E., submucosal edema.

In summary, we could show that a more distantly related competitor does not compete as well in the gut but nonetheless is able to abolish systemic invasion and intestinal inflammation, and this may represent a more easily translatable model. Moreover, future analysis of the metabolism of *S*.Tm and commensal *E. coli* strains may allow for the identification of strain combinations with a more complete metabolic niche overlap.

Taken together, we here demonstrate that the combination of an inactivated oral vaccine and a competitor strain in the gut lumen can protect from *S*.Tm colonization and disease in murine models. The extent of gut lumen clearance was well-correlated with fecal lipocalin-2 and intestinal inflammation.

### Comparing vaccination/niche competition to licensed animal vaccines reveals mechanistic differences in protection of the intestine and systemic sites

Live-attenuated non-typhoidal *Salmonella* vaccines, carrying a mutation in *aroA*, which renders the strain auxotrophic for aromatic amino acids, are already used for broilers^30^ and very similar auxotrophic strains are used in pig-farming^31^. In order to benchmark our approach to a known *S*.Tm vaccine we therefore compared protective efficacy and safety of the combined inactivated oral vaccine+*Ec*^8178^ treatment to that of *S*.Tm^aroA^ vaccination. To gain mechanistic insight into protection at the tissue level versus the gut lumen, we also compared protection in the NTS model to protection in the murine oral typhoid model, in which no major gut luminal niche is generated^32^.

In the NTS model (**Fig. 5A**), both the classical *S*.Tm^aroA^ vaccination as well as EvoTrap vaccination+*Ec*^8178^ led to the induction of *S*.Tm^WT^ specific antibodies (**Fig. 5B**) and prevented weight loss (**Fig. 5C**). EvoTrap vaccination+*Ec*^8178^ resulted in significantly lower levels of *S*.Tm^WT^ in feces and cecum (**Fig. 5D and E**) and completely prevented intestinal inflammation (**Fig. 5F-H**). *S*.Tm^aroA^ vaccination, in contrast, showed no effect on fecal or cecal *S*.Tm loads (**Fig. 5D and E**) and only a very small suppression of intestinal inflammation (**Fig. 5F-H**). Interestingly, although EvoTrap vaccination+*Ec*^8178^ showed better protection at the level of the gut and mesenteric lymph nodes (**Fig. 5D, E and I**), the classical *S*.Tm^aroA^ vaccination provided better protection from systemic *S*.Tm^WT^ infection. Liver and spleen CFU of *S*.Tm^WT^ were not dramatically decreased in this experiment by EvoTrap vaccination+Ec^8178^ but were strongly reduced by vaccination with live *S*.Tm^aroA^ (**Fig. 5J and K**). Interestingly, although *S*.Tm^aroA^ was absent from cecum and feces it could still be found in MLNs and liver at day 46 post-vaccination (**Fig. 5I-K**), consistent with known safety challenges associated with these live-attenuated vaccines^6^.

**Figure 5.**
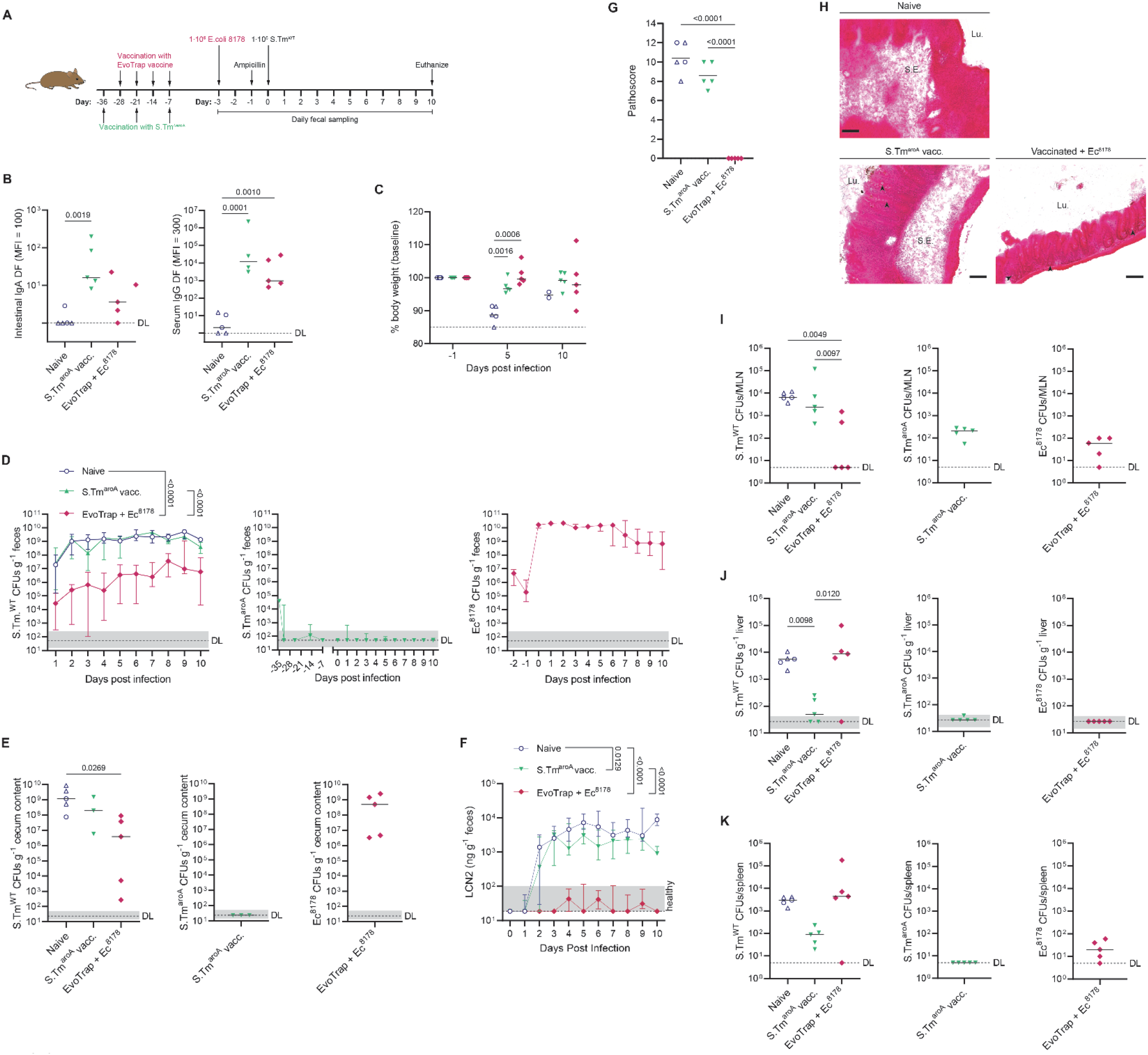
EvoTrap vaccination together with niche competition provides better intestinal protection than a licensed animal vaccine in the murine NTS model. PBS (blue circles), EvoTrap-vaccinated (pink diamonds) or *S*.Tm^aroA^ vaccinated (green triangles) 129S6/SvEv mice were pretreated with ampicillin and infected with 1·10^6^ *S*.Tm^WT^. EvoTrpa-vaccinated mice were pre-colonized with 1·10^9^ *E. coli* 8178 3 days before infection. (**A**) Experimental procedure. (**B**) *S*.Tm^WT^-specific intestinal IgA and serum IgG titres as determined by flow cytometry. (**C**) Change in body weight over the course of infection. Solid lines show the mean. Fecal (**D**) and cecum content (**E**) CFUs as determined by selective plating. Intestinal inflammation as determined by fecal lipocalin-2 (**F**) and histopathological scoring of cecal tissue sections (**G**). Solid lines in (G) show the mean. (**H**) Representative images of H&E-stained cecal tissue sections. Arrowheads show exemplary goblet cells. Scale bars: 100 µm. (**I-K**) CFUs in MLN (**I**), liver (**J**) and spleen (**K**). N = 5 mice/group. Solid lines depict the median unless stated otherwise, error bars the interquartile range. Dotted lines show the detection limit and the shaded area the range for cases in which the detection limit is dependent on sample weight. Open triangles show mice that had to be euthanized prematurely due to excessive weight loss (≥ 15%) or disease symptoms. Statistics were performed by mixed-effects analysis (C) or one-way ANOVA (C, G) on log-normalized data (B, E, I-K) or area under the curve (AUC) (D, F). CFU, colony forming unit; DF, dilution factor; LCN2, lipocalin-2; Lu., lumen; MFI, median fluorescence intensity; MLN, mesenteric lymph node; S.E., submucosal edema.

In the murine typhoid model **(Fig. 6A**), both vaccine regimens could prevent intestinal colonization (**Fig. 6B and C**) and supress any detectable increase in intestinal lipocalin-2 levels (**Fig. 6D**). MLN colonization (**Fig. 6E**) correlated well with intestinal colonization levels at day 10 post infection. As in the murine NTS model, vaccination with *S*.Tm^aroA^ resulted in superior protection against infection of liver and spleen (**Fig. 6F and G**). However, persistent *S*.Tm^aroA^ was again found in systemic sites a full 17 days after the last vaccination.

**Figure 6.**
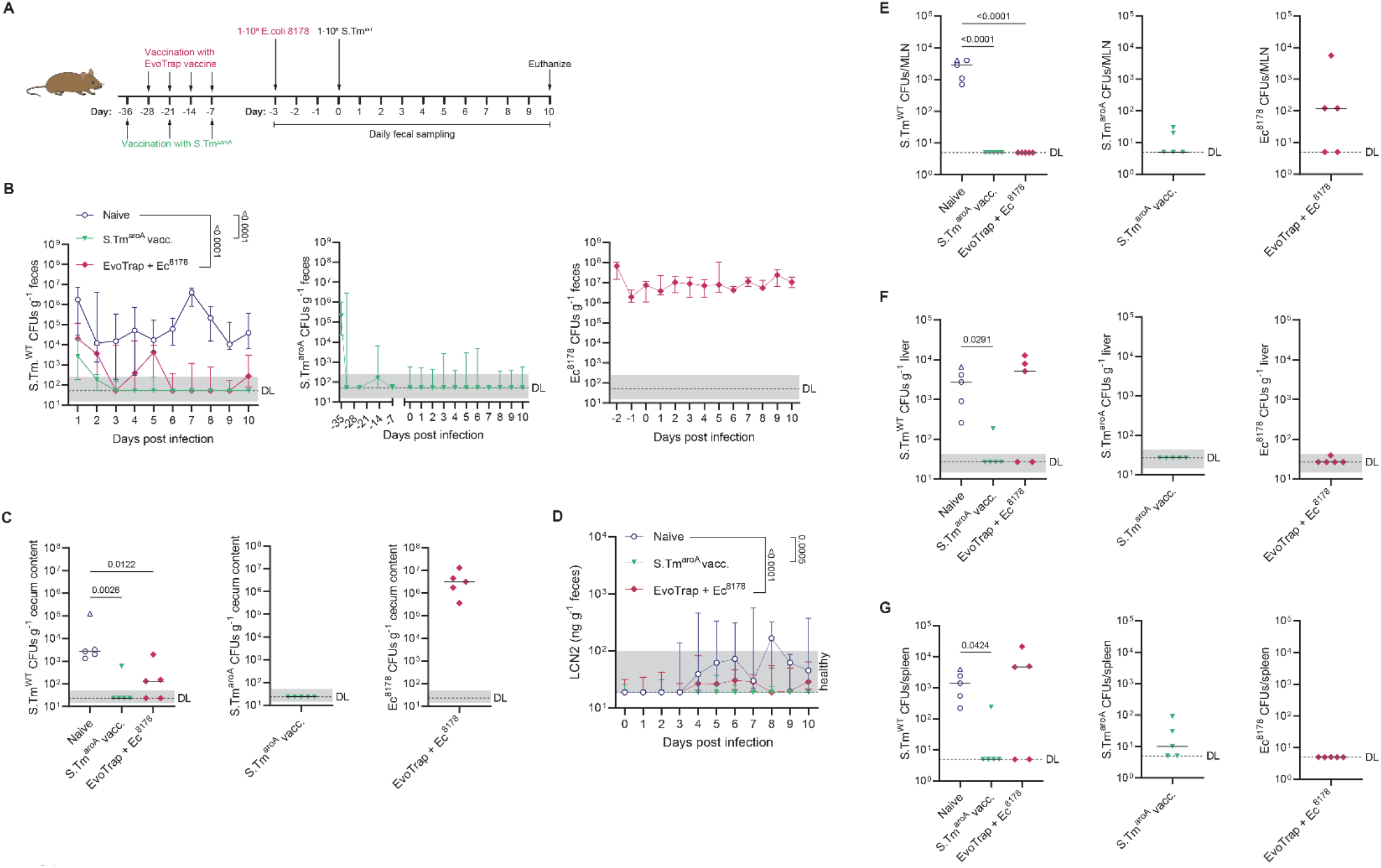
*S*.Tm^aroA^ vaccination protects better against systemic invasion of *S*.Tm^WT^ in the murine Thyphi model. PBS (blue circles), EvoTrap-vaccinated (pink diamonds) or *S*.Tm^aroA^ vaccinated (green triangles) 129S6/SvEv mice were infected with 1·10^6^ *S*.Tm^WT^ without antibiotic pretreatment. EvoTrpa-vaccinated mice were pre-colonized with 1·10^9^ *E. coli* 8178 3 days before infection. (**A**) Experimental procedure. Fecal (B) and cecum content (**C**) CFUs as determined by selective plating. (**D**) Intestinal inflammation as determined by fecal lipocalin-2. (**E-G**) CFUs in MLN (**E**), liver (**F**) and spleen (**G**). N = 5 mice/group. Solid lines depict the median, error bars the interquartile range. Dotted lines show the detection limit and the shaded area the range for cases in which the detection limit is dependent on sample weight. Open triangles show mice that had to be euthanized prematurely due to excessive weight loss (≥ 15%) or disease symptoms. Statistics were performed by ANOVA on log-normalized data (C, E-G) or area under the curve (AUC) (B, D). CFU, colony forming unit; LCN2, lipocalin-2; MLN, mesenteric lymph node.

These data imply that protection of the gut tissue and of systemic sites require different mechanisms and can be uncoupled: vaccination/niche competition provides superior protection of all examined gut sites and gut-draining lymphoid tissues, while live-attenuated vaccines provide better protection of deep systemic sites. This therefore reveals both a promising approach for improved safe prophylaxis of intestinal bacterial infections and a system to investigate missing stimuli from whole-cell inactivated oral vaccines to improve their protection of systemic sites.

## Discussion

Previous work from our group showed the importance of IgA in pathogen immune exclusion and provided a mechanistic explanation for this phenotype based on IgA-driven enchained growth^16^. However, the extent of tissue protection that can be provided by IgA alone is typically too weak to completely protect intestinal tissues. A further feature of secretory IgA is its ability to increase the clearance rate of bacteria from the gut lumen^16^, but in the case of a large open niche this function affects evolutionary dynamics without affecting total population size. In contrast, if a niche competitor is present and either replicates faster, or is cleared more slowly, or ideally both, this competitor has a fitness advantage and can rapidly take over the niche. Exploiting IgA to increase the clearance rate of the pathogen allows fast and efficient niche competition, generating high-efficacy prophylaxis with a very safe inactivated whole-cell oral vaccine.

This observation may explain some controversies in the existing *Salmonella* literature. For example, the extent of protection from NTS obtained with live-attenuated vaccines in different laboratories varies extensively^33^. Our data indicate that the efficacy of *Salmonella* niche competitors in the microbiota of mice, which likely varies in SPF facilities around the world, is a critical determinant of protective efficacy. Moreover, this is generally in line with the concept of colonization resistance^34^: the microbiome itself provides extensive protection from *Salmonella* gut colonization, in part due to direct competition for metabolic resources. It makes complete sense that intestinal antibody responses have evolved to work with this protective effect of the microbiota.

A major question remaining is how to ideally select niche competitors that are genuinely benign but also highly effective. Niche competition will occur most extensively when the environmental and metabolic requirements of the competitor strain are as close as possible to the pathogen. Using modified versions of the pathogen itself provides the ideal candidate as by definition this must compete for an identical niche. However, genetically modified pathogens may be a very hard sell as “probiotics”. An alternative would be to make use of commensal bacterial isolates that are rigorously checked for potential pathogenicity. *E. coli* strains, such as Nissle 1917 are of course attractive propositions for this both due to proven safety^35^ and relatively short genetic distances from the pathogen. However, non-typhoidal *Salmonella* are metabolic generalists, and it remains possible that there is no one ideal competitor for all microbiota perturbations. Using a bacterial consortium instead of only one strain as competitor might be able to circumvent this problem by using bacterial strains that complement each other. A mix of three E. coli strains has shown increased protection against *S*.Tm infection after dietary perturbation when compared to only one of those E. coli strains^29^. We clearly demonstrate that a murine commensal *E. coli* 8178 is less effective in clearing *Salmonella* from the gut lumen than a *Salmonella*-derived competitor. Nevertheless, this is sufficient to prevent intestinal inflammation over a long course of infection. An adaption in the administration of *E. coli* 8178 to a more realistic setting i.e. giving it without prior antibiotic pre-treatment, led to a dramatic decrease in protection of systemic sites from *S*.Tm^WT^. We can currently only speculate regarding the underlying mechanism of protection by high-density *E. coli* colonization. As *E. coli* overgrowth is generally not considered to be part of a healthy microbiota, this is not promising for translation. However, we have a few clues: We do not assume that this is due to direct competition in these sites as we could not detect *E. coli* 8178 in spleen or liver of vaccinated mice in case of antibiotic pre-treatment. An alternative would be *E. coli-*induced changes in intestinal innate immune signalling^36^ with either direct effects on *Salmonella* invasion, or indirect effects via changes in abundance of factors such as bile acids^37^ or short-chain fatty acids^38,39^ that repress major virulence factor expression. Another concern is potential immunogenicity of the inoculated niche competitors over time, which could decrease their efficacy or lead to clearance before they are needed, as well as waning of the intestinal IgA response over time. Novel techniques to identify robust niche competitor strains will be needed in order to translate the approach into human use, as well as further studies on long-term robustness of protection. Within veterinary applications, we would hope that sterilizing-immunity could be generated acutely across whole farms resulting in permanent de-colonization of the herds, and potentially circumventing any potential problems with response longevity.

Due to improved efficacy, the field of non-typhoidal *Salmonella* vaccine research has been strongly focused on live-attenuated vaccines. *S*.Tm vaccines are licensed only for the veterinary sector and both the poultry and swine vaccine are live, auxotrophic *S*.Tm strains^30,31^. These vaccine strains were originally tested in mice and showed protection against subsequent lethal infection with wildtype *S*.Tm^7^. Although *S*.Tm^aroA^ strains were stated to be replication incompetent and avirulent, later studies with these mutants showed repeatedly higher bacterial counts than the applied infection doses, long-lasting persistence in liver and spleen and severe splenomegaly^40,41^. Continuous shedding of a *S*.Tm *aroA ssaV* double mutant strain was also observed in a small study with healthy human volunteers for up to 23 days^6^ and a *S*.Tm *phoP/phoQ* deletion strain although in this study they treated volunteers still shedding the vaccine strain with antibiotics from day nine onwards^42^. Moreover, while immunocompetent mice do survive infection with *S*.Tm^aroA^, mice lacking CD4+ T cells or IFNγ succumb to this type of vaccination^43,44^. However, individuals with immunodeficiencies or a not yet fully developed immune system are the demographic group most requiring effective *Salmonella* prophylaxis, and it is therefore vital to develop a vaccine that is also safe to use in high-risk individuals.

Because of this we have directly compared protective efficacy of the *aroA* live-attenuated vaccine and the combination of an “Evolutionary Trap” version of the inactivated whole-cell oral vaccine with an *E. coli* niche competitor. Our data clearly demonstrate a much stronger protection of the intestinal environment by the inactivated vaccine/niche competitor approach, indicating that combined secretory IgA and competition robustly prevent invasion of intestinal tissues and intestinal inflammation. In contrast, the live-attenuated vaccines provided superior protection of systemic sites. As both vaccines induce broad and robust IgG responses, we could exclude this mechanism as the grounds for protection. Instead, there are two potential explanations: 1) the live-attenuated vaccine permanently colonizes systemic sites, generating low-grade inflammation and niche competition in these environment (i.e. protection independent of adaptive immunity) or 2) live-attenuated vaccines more efficiently induce cell-mediated immunity which can control the intracellular stages of *Salmonella* infection. We cannot currently distinguish between these phenomena based on our data, but note that CFU of live *S*.Tm^aroA^ could be detected in the spleen and liver of vaccinated mice more than two weeks after the final vaccination which would be consistent with explanation (1). Nevertheless, live-attenuated vaccines are also expected to induce stronger systemic T cell responses than inactivated bacterial cells^45^. Overall, our conclusion is that vaccination/niche competition is a promising approach to prevent intestinal disease and its transmission using a very safe vaccine. There is still room for improvement in preventing Salmonella invasion and replication at systemic sites. However, it possible that this strategy can achieve full protection in more realistic infection scenarios with less microbiota disruption, reduced niche sizes for pathogen blooms and lower inoculum sizes.

Overall, this work reveals the ability of rationally designed oral vaccines combined with carefully selected niche competitors to prevent or supress intestinal colonization by a pathogen. Even in the very stringent non-typhoidal Salmonellosis model, this strategy is capable of generating a type of “sterilizing immunity”, i.e. complete clearance of the pathogen, in the gut lumen within one week of challenge. This is also highly suggestive of a role of secretory IgA in controlling competition between intestinal microbiota members, which is consistent with observations of instability of *E. coli* at the strain level, but not the species level, over time in healthy volunteers^46^. Therefore, this approach also has potential applications in microbiota engineering where strain replacement could either eliminate a detrimental strain or could introduce a strain with beneficial properties into an otherwise complete microbiota.

## Materials and Methods

All materials are available upon request to the corresponding authors.

### Strains and plasmids

All strains and plasmids used in this study are listed **Table S1**.

Bacteria were cultivated in lysogeny broth (LB) containing appropriate antibiotics (100 µg/ml streptomycin (AppliChem); 15 µg/ml chloramphenicol (AppliChem); 50 µg/ml kanamycin (AppliChem); 50 µg/ml ampicillin (AppliChem)). Dilutions were prepared in Phosphate Buffered Saline (PBS, Difco).

Gene-deletion mutants were created by generalized transduction with bacteriophage P22 HT105/1 *int-201* as described in^47^. When needed, antibiotic resistance cassettes were removed using the temperature-inducible FLP recombinase encoded on pCP20^48^. Deletions originated from in-frame deletions made in *S*.Tm 14028S, kind gifts from Prof. Michael McClelland (University of California, Irvine). Primers used for verifications of gene deletions or genetic background are listed **Table S2**.

Plasmids were transferred by electro-transformation into competent cells^29,49^.

### Mice

All animal experiments were performed in accordance with Swiss Federal regulations approved by the Commission for Animal Experimentation of the Kanton Zurich (licenses 193/2016, 158/2019 and 120/2019; Kantonales Veterinäramt Zürich, Switzerland). Specific opportunistic pathogen-free (SPF, containing a complete microbiota free of an extended list of opportunistic pathogens) 129S6/SvEvTac WT mice were used in all experiments except for Fig. S1 A-F, where C57BL/6J WT mice were used. Mice were bred and housed in individually ventilated cages with a 12 h light/dark cycle in the ETH Phenomics Center (EPIC, RCHCI), ETH Zürich and were fed a standard chow diet. All mice included in experiments were 4 weeks or older and objectively healthy as determined by routine health checks. Wherever possible an equal number of males and females was used in each group. Mice were allocated cage-wise to groups with a minimum of two cages per group. As strong phenotypes were expected, we adhered to standard practice of analysing at least five mice per group. Researchers were not blinded to group allocation to decrease the risk of contamination and because the majority of readouts were quantitative and not subjective (CFU determination, ELISA). The one exception to this was histopathology scoring, for which a blinded researcher (SAF) not otherwise involved in the experiment carried out all scoring.

### Vaccinations

Mice were either vaccinated with peracetic acid (PA) killed vaccines or live-attenuated *S*.Tm^aroA^.

Peracetic acid killed vaccines were produced as previously described^50^. Briefly, bacteria were grown overnight to late stationary phase, harvested by centrifugation and re-suspended to a density of 10^9^-10^10^ per ml in sterile PBS. Peracetic acid (Sigma-Aldrich) was added to a final concentration of 0.4% v/v. The suspension was mixed thoroughly and incubated for 60 min at room temperature. Bacteria were washed three times in 50-100 ml sterile PBS. The final pellet was resuspended to yield a density of 10^11^-10^12^ particles per ml in sterile PBS. The exact number was determined by flow cytometry with counting beads (Fluoresbrite® Multifluorescent Microspheres). Vaccines were stored at 4 °C for up to three weeks. Each batch of vaccine was tested for sterility before use. Vaccine lots were released for use only when a negative enrichment culture had been confirmed. Mice were vaccinated with 10^10^-10^11^ PA-killed bacteria by oral gavage, once weekly for 4 weeks. Where multiple strains were used, equal numbers of each strain were given.

Live-attenuated *S*.Tm^aroA^ was grown overnight in LB containing chloramphenicol. The cells were washed in PBD and resuspended at a density of 10^10^ bacteria per ml. Mice were orally vaccinated with 10^9^ *S*.Tm^aroA^ in 100 μl three time in bi-weekly intervals without antibiotic treatment.

Vaccinations were started in mice at an age of 4-6 weeks.

### Colonization with the bacterial niche competitor and *Salmonella* challenge infections

The competitor strain was grown overnight in LB containing the appropriate antibiotics. In the morning, the bacteria were washed with sterile PBS and diluted. The competitor was introduced by oral gavage into the respective groups either at 5·10^3^ CFUs after antibiotic pre-treatment or at 1·10^9^ CFUs without antibiotic pr-treatment of the animals.

Non-typhoidal *Salmonella* infections were carried out as previously described^21^. In brief, mice were orally pretreated 24 h before infection with 25 mg streptomycin or 20 mg ampicillin. Strains were cultivated overnight separately in LB containing the appropriate antibiotics. Subcultures were prepared before infections by diluting overnight cultures 1:20 in fresh LB without antibiotics and incubation for 3 h at 37 °C. The cells were washed in PBS, diluted, and 100 µl of bacteria were used to infect mice by orogastric gavage with either 5·10^3^ or 1·10^6^ *S*.Tm CFUs, as indicated in the respective figure legends/text. Competitions were performed by inoculating 1:1 mixtures of each competitor strain. For mouse typhoid-like infection, the animals were infected with 1·10^6^ *S*.Tm CFUs without prior antibiotic treatment. A detailed layout of the vaccination and infection schedule is shown in the figures.

Feces were sampled daily, homogenized in 500 μl PBS by bead beating (3 mm steel ball, 25 Hz for 2.5 min in a TissueLyser (Qiagen)), and large particles were sedimented by centrifugation at 500x *g* for 1 minute. Bacteria were enumerated by selective plating on MacConkey agar supplemented with the appropriate antibiotics. Fecal samples for lipocalin-2 measurements were kept homogenized in PBS at -20 °C. At endpoint, blood was collected from the heart into 1.1 ml serum gel tubes (Sarstedt). Intestinal lavages were harvested by flushing the small intestinal content with 2 ml of PBS using a cannula. The middle part of the cecum was placed into OCT Compound (Tissue-Tek), snap-frozen and stored at -80 °C until analysis. Spleen, liver, mesenteric lymph nodes were collected and homogenized in 1 ml PBS at 30 Hz for 3 min. Cecum content was collected and homogenized in 500 μl PBS at 25 Hz for 2.5 min. After centrifugation at 500x *g* for 1 minute, bacteria were plated on selective MacConkey agar.

### Quantification of fecal lipocalin-2

Fecal pellets were processed as described above. Homogenized feces was centrifuged at 16000x *g* for 5 min and the resulting supernatant was analysed in duplicates using the mouse lipocalin-2 ELISA duoset (R&D, DY1857) according to the manufacturer’s instructions.

### Analysis of specific antibody titres by bacterial flow cytometry

Specific antibody titres in mouse intestinal washes and serum were measured by flow cytometry as described^51^. Briefly, intestinal washes and blood were collected as described above. Blood was centrifuged at 10000x *g* for 5 min to obtain serum, heat-inactivated at 56 °C for 30 min and stored at -20 °C until further analysis. Intestinal lavage was centrifuged at 16000x *g* for 5 min to clear all bacterial-sized particles and stored at -20 °C until analysis. Bacterial targets (antigen against which antibodies are to be titred) were grown overnight in LB, then gently pelleted for 2 min at 7000x *g*. The pellet was washed with 0.2 µm-filtered PBS before resuspending at a density of approximately 10^7^ bacteria per ml. After thawing, intestinal washes were centrifuged again at 16000x *g* for 5 min. Supernatants were used to perform serial dilutions. 50 μl of the dilutions were incubated with 50 μl bacterial suspension for 15 min at room temperature. Bacteria were washed twice with 150 μl PBS by centrifugation at 7000x *g* for 5 min, before resuspending in 25 μl of 0.2 µm-filtered PBS containing polyclonal Alexa Fluor 647 Rabbit Anti-Mouse IgG (Jackson ImmunoResearch, 15 µg/ml, 315-605-003, AB_2340239) or monoclonal Brilliant Violet 421 Rat Anti-Mouse IgA (BD Bioscience, 2 µg/ml, 743293, AB_2741405). After 5 min of incubation at RT, bacteria were washed twice with PBS as above and resuspended in 100 μl PBS for acquisition on a Beckman Coulter Cytoflex S using FSC and SSC parameters to threshold acquisition in logarithmic mode. Data were analysed using FlowJo (Treestar). After gating on bacterial particles, log-median fluorescence intensities (MFI) were plotted against lavage dilution factor for each sample and 4-parameter logistic curves were fitted using Prism (Graphpad, USA). Titers were calculated from these curves as the dilution factor giving an above-background signal (typically MFI=300).

### Histological procedures

Tissue embedded in OCT Compound was cut into 5 μm cryosections and mounted on glass slides. Cryosections were air dried overnight at room temperature and stained with hematoxylin and eosin (H&E). Scoring of cecal inflammation was done in a blinded manner assessing the following four criteria as previously described^21^.

i. Submucosal edema. Submucosal edema was scored as follows: 0 = no pathological changes; 1 = mild edema (the submucosa accounts for <50% of the diameter of the entire intestinal wall - tunica muscularis to epithelium); 2 = moderate edema (the submucosa accounts for 50 to 80% of the diameter of the entire intestinal wall); and 3 = profound edema (the submucosa accounts for >80% of the diameter of the entire intestinal wall).
ii. PMN infiltration into the lamina propria. Polymorphonuclear granulocytes (PMN) in the lamina propria were enumerated in 10 high-power fields (x400 magnification; field diameter of 420 μm), and the average number of PMN/high-power field was calculated. The scores were defined as follows: 0 = <5 PMN/high-power field; 1 = 5 to 20 PMN/high-power field; 2 = 21 to 60/high-power field; 3 = 61 to 100/high-power field; and 4 = >100/high-power field. Transmigration of PMN into the intestinal lumen was consistently observed when the number of PMN was >60 PMN/high-power field.
iii. Goblet cells. The average number of goblet cells per high-power field (magnification, x400) was determined from 10 different regions of the cecal epithelium. Scoring was as follows: 0 = >28 goblet cells/high-power field; 1 = 11 to 28 goblet cells/high-power field; 2 = 1 to 10 goblet cells/high-power field; and 3 = <1 goblet cell/high-power field.
iv. Epithelial integrity. Epithelial integrity was scored as follows: 0 = no pathological changes detectable in 10 high-power fields (x400 magnification); 1 = epithelial desquamation; 2 = erosion of the epithelial surface (gaps of 1 to 10 epithelial cells/lesion); and 3 = epithelial ulceration (gaps of >10 epithelial cells/lesion; at this stage, there is generally granulation tissue below the epithelium). The combined pathological score for each tissue sample was determined as the sum of these scores. It ranges between 0 and 13 arbitrary units and covers the following levels of inflammation: 0 = intestine intact without any signs of inflammation; 1 to 2 = minimal signs of inflammation (this was frequently found in the ceca of SPF mice; this level of inflammation is generally not considered as a sign of disease); 3 to 4 = slight inflammation; 5 to 8 = moderate inflammation; and 9 to13 = profound inflammation.

### Statistical analysis

Sample size was determined based on previous experiments^20^. Where large effect sizes where expected a minimum of five mice/group were used. Researchers were not blinded for the assignment of the experiments and the data analysis except for histopathological scoring. Where errors are expected to be log-normal distributed (e.g., CFU determination or ELISA data based on serial dilutions), all statistical tests were carried out on log normalized data. Where two groups of data were compared, analysis was carried out using unpaired two-tailed t-test. One-way ANOVA followed by Tukey’s Test was used for comparison of three or more groups. Statistical analysis on time course data was either done by mixed-effects analysis or by calculating the area under the curve (AUC) of the log normalized data and then assessing differences in AUC with one-way ANOVA and Tukey’s test. Statistical analysis was performed with Graphpad Prism Version 9.2.0 for Windows (GraphPad Software, La Jolla, California USA). P values of less than 0.05 were reported.

## Acknowledgements

We thank Daniel Hoces and Markus Arnoldini, and other members of the Slack lab for helpful discussions and comments, and the staff at the RCHCI and EPIC animal facilities for their excellent support.

Funding for this work was provided by the Gebert Rüf Microbials (GR073_17). VL, AW, CL and ES are supported by the Gebert Rüf Microbials (GR073_17). ES acknowledges the support of the Swiss National Science Foundation (40B2-0_180953, 310030_185128), and European Research Council Consolidator Grant (865730). This work was supported as a part of NCCR Microbiomes, a National Centre of Competence in Research, funded by the Swiss National Science Foundation (grant number 180575). Funding was provided by the Botnar Research Centre for Child Health as part of the Multi-Investigator Project: Microbiota Engineering for Child Health. MD is supported by a SNF professorship (PP00PP_176954) and Gebert Rüf Microbials (PhagoVax GRS-093/20). WDH acknowledges funding by grants from the Swiss National Science Foundation (310030_192567, NCCR Microbiomes). CL is supported by Agence Nationale de la Recherche (ANR-21-CE45-0015, 376 ANR-20-CE30-0001) and MITI CNRS AAP adaptation du vivant à son environnement.

## Author contributions

Verena Lentsch, Conceptualization, Investigation, Methodology, Validation, Data Curation, Project administration, Formal Analysis, Visualization, Supervision, Writing – original draft, Writing – review and editing; Aurore Woller, Data curation, Formal analysis, Investigation, Methodology, Resources, Software, Validation, Visualization, Writing – review and editing; Claudia Moresi, Investigation, Validation, Data Curation, Writing – review and editing; Stefan A. Fattinger, Investigation, Writing – review and editing; Selma Aslani, Investigation, Writing – review and editing; Wolf Dietrich Hardt, Resources, Supervision, Writing – review and editing; Claude Loverdo, Formal analysis, Methodology, Resources, Software, Validation, Writing – review and editing; Médéric Diard, Conceptualization, Writing – review and editing; Emma Slack, Conceptualization, Funding acquisition, Project administration, Resources, Supervision, Writing – original draft, Writing – review and editing.

## Supplementary Materials and Methods

### Estimation of *in vivo* growth rates in the gut by plasmid dilution

Absolute *S*.Tm growth rates in the gut were assessed using replication-incompetent plasmid pAM34, which has been described previously^29,49^. Briefly, pAM34 is a ColE1-like vector in which the replication of the plasmid is under the control of the LacI repressor, whereby plasmid replication only occurs in the presence of isopropyl β-d-1-thiogalactopyranoside (IPTG). *S*.Tm carrying the pAM34 plasmid was therefore cultured overnight in the presence of 1 mM IPTG in LB containing streptomycin. Cultures were diluted 1:20 into fresh LB broth without IPTG or antibiotics and sub-cultured for 3 h at 37 °C. Inocula for infection were prepared as described above. Concurrently, the inoculum was serially diluted into fresh LB broth without IPTG and cultured for 20 h at 37 °C to generate a standard curve relating plasmid loss to generations undergone for each experiment. pAM34-carrying bacteria within the overnight cultures and the feces were determined by selective plating on agar plates containing 50 µg/ml ampicillin and 1 mM IPTG. To quantify the total population size, samples were further plated on agar plates containing 100 µg/ml streptomycin. The fraction of pAM34-carrying bacteria was calculated using the ratio of pAM34-carrying CFU to the total population CFU and generations estimated by interpolation from the matched standard curve.

### Non-typhoidal *Salmonella* transmission

Donor mice were vaccinated with PA-*S*.Tm as described above, orally pretreated with 25 mg streptomycin, and colonized 24 h later with 5·10^3^ *S*. Tm^Comp^. 2 days later, mice were treated again with 25 mg Streptomycin by orogastric gavage, and 24 h later infected with 1·10^6^ *S*.Tm^WT^. On day 9 post infection, one fecal pellet was collected from each mouse, weighed, and homogenized in 200 μl PBS. Large debris was removed by centrifugation at 500x *g* for 1 minute and 50 μl of the supernatant were immediately given by oral gavage to streptomycin pretreated naïve recipient mice. As a control, the same procedure was done using naïve mice without competitor colonization as donor mice. Recipient mice were euthanized, and organs were collected on day 3 post transmission. In both donor and recipient mice, fecal pellets were collected daily and selective plating was used to enumerate *Salmonella* and determine the relative proportions of both competing bacterial strains.

### Flow cytometry for analysis of O:5 and O:12-0 intensity on *Salmonella* clonal cultures

Overnight cultures (1 µl) made in 0.2 µm-filtered lysogeny broth was stained with 0.2 µm-filtered solutions of STA5 (human recombinant monoclonal IgG2 anti-O:12-0, 3.2 µg/ml)^16^ or rabbit anti-Salmonella O:5 (Difco, 1:200, 226601). After incubation at 4 °C for 30 min, the bacteria were washed twice by centrifugation at 7000x *g* and resuspension in PBS/2% BSA. The bacteria were then resuspended in 0.2 µm-filtered solutions of appropriate secondary reagents (Alexa 647-anti-human IgG (Jackson ImmunoResearch, 1:100, 109-605-098, AB_2337889) and Brilliant Violet 421-anti-rabbit IgG (BioLegend, 1:100, 406410, AB_10897810)). This was incubated for 30 min at 4 °C before the cells were washed as above and resuspended for acquisition on a Beckman Coulter Cytoflex S.

### LPS purification and silver staining

LPS was isolated by applying the hot phenol–water method^52^, followed by buffer exchange against 15 ml PBS and concentration in 500 μl PBS. LPS samples were separated on a 13% Tricin gel by gel electrophoresis and silver staining was performed as described previously^53^.

### Mathematical modeling of *S*.Tm^WT^ extinction in the gut

All mathematical modelling is based on the arithmetic mean of the experimental data.

For Eq. 1 we consider a case where K_2_ ≪ K_1_ because the height of final *S*.Tm^Comp^ “plateau” observed in competition data (which is proportional to K_2_, see calculations below) is much lower than the size of *S*.Tm^Comp^ population at day 1 (which is proportional to K_1_, see calculations below). We successively consider the cases K_1_ ≪ K_3_ and K_3_ ≪ K_1_ and show that the latter is more compatible with the competition data. In the competition data (vaccinated group with *S*.Tm^Comp^), several distinct regimes can be observed (**Fig. S6**), which is also highlighted below with the model.

Case where K_3_ ≫ K_1_

Regime R_1_ with W, M, C ≪ K_1_

In this case, the equations become

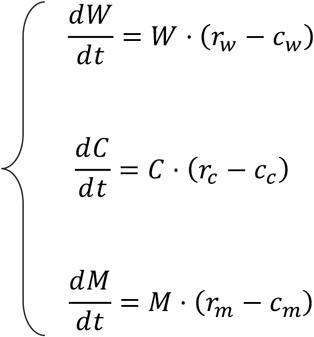

Thus, we have

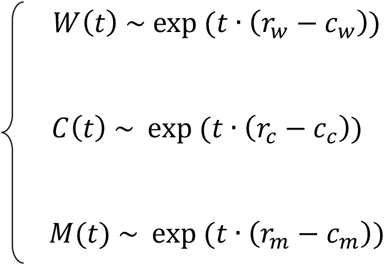

Regime R_2_ with W, M ≪ C∼ Θ(K_1_)

In this case, the equations become

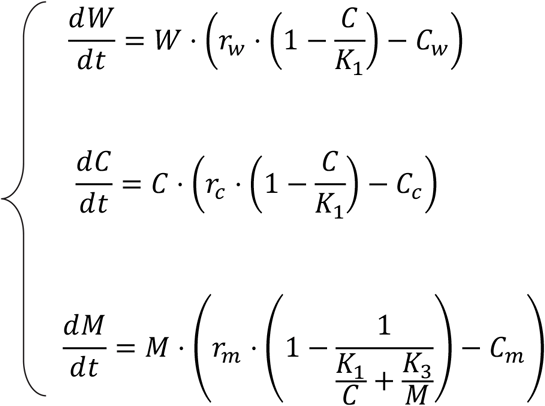

Note: in this regime, 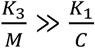. Thus, we have

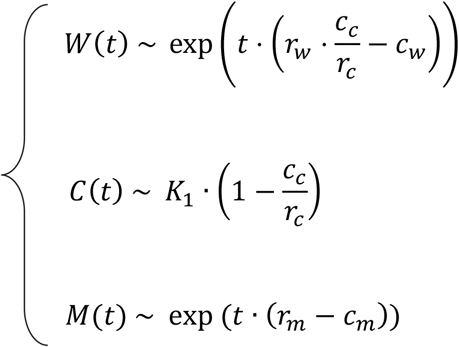

Note that we assume C is the first to be close to the carrying capacity K_1_. Next, at t = τ, we have M ∼ C, that is

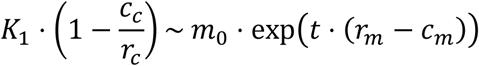

Regime R_3_ with C, W ≪ M and C ≫ K_2_

In this regime, we have 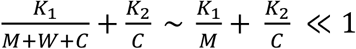. This implies that there are no nutrients available to C and W. In this case, the equations become

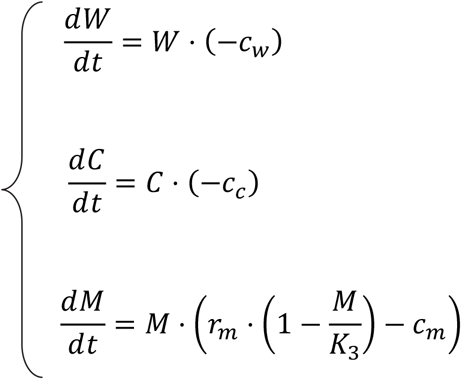

Thus, we have

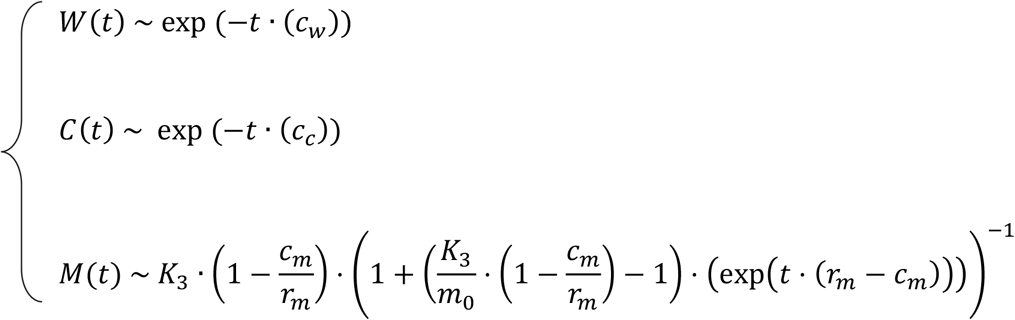

Thus, in this regime, the slopes expected for log (C) and log (W) are -c_c_ and -c_w_, respectively. Thus, if we estimate the slopes of log (C) and log (W) from *in vivo* data (vaccinated group) for R_3_, we should get an estimation of the clearance rate values. Table S3 shows the slopes obtained from experiments: they range from -1.8 to -3.9, which would correspond to extremely slow clearance rates. This suggests that the case K_1_ ≫ K_3_ is probably not well suited to describe the data.

**Table S3:** Estimation of the slopes of log (C) and log (W) from *in vivo* data (vaccinated group + *S*.Tm^Comp^). The estimation was performed for different possible durations of this regime.

| Width of the regime | Slope for log (C) | Slope for log (W) |
| --- | --- | --- |
| day 3 to day 7 | -3.4 day<sup>-1</sup> | -2.2 day<sup>-1</sup> |
| day 3 to day 6 | -3.7 day<sup>-1</sup> | -1.8 day<sup>-1</sup> |
| day 4 to day 7 | -3.3 day<sup>-1</sup> | -2.4 day<sup>-1</sup> |
| day 4 to day 6 | -3.9 day<sup>-1</sup> | -1.9 day<sup>-1</sup> |

Regime R_4_ with W, M ≪ C∼ Θ(K_2_)

In this case, the equations become

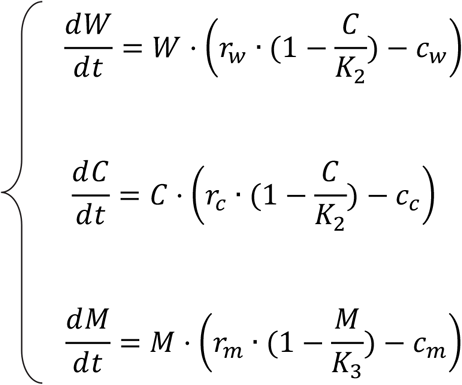

Thus, at long times we have

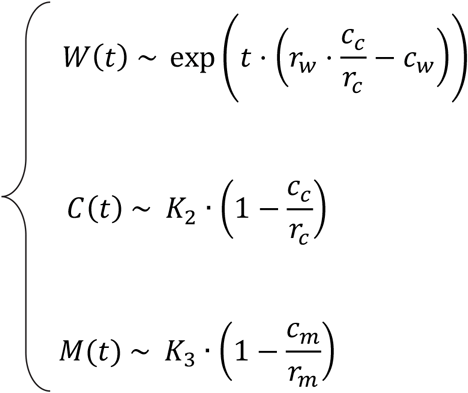

Case where K_1_ ≫ K_3_

Regime R_1_ with W, M, C ≪ K_1_

If we also have M ≪ K_3_, the equations become

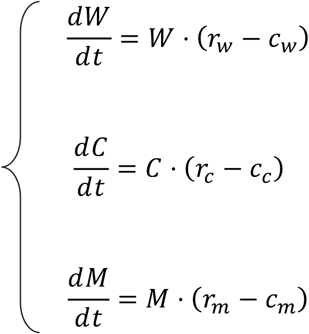

Thus, we have

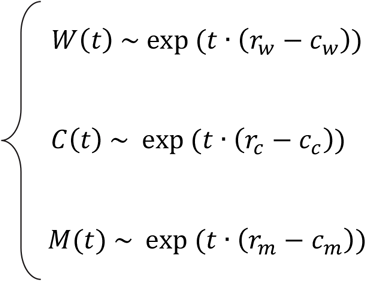

Regime R_2_ with W, M ≪ C∼ Θ(K_1_)

In this case, the equations become

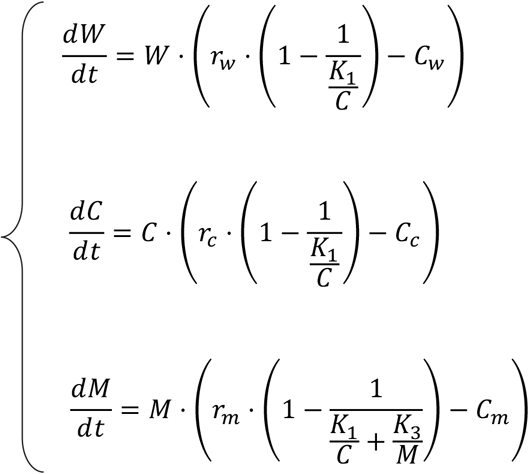

Thus, we have

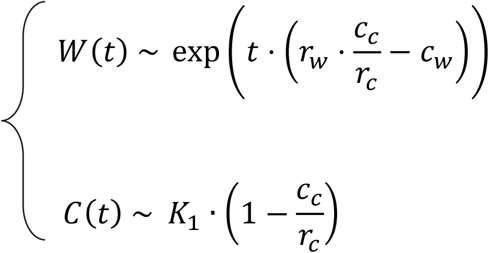

When M is in a regime such that K_3_ ≪ M ≪ K_1_, then 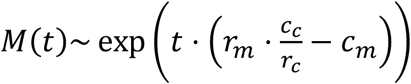.

Regime R_3_ with C, W ≪ M and C ≫ K_2_

In this case, the equations become

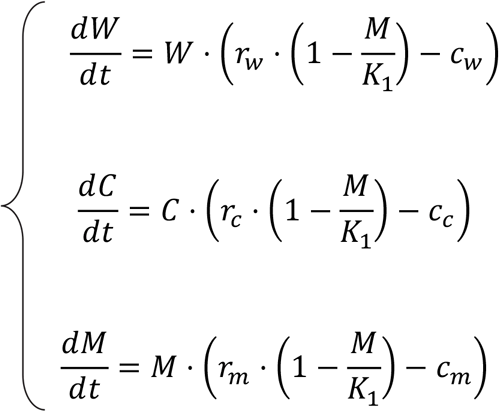

Thus, we have

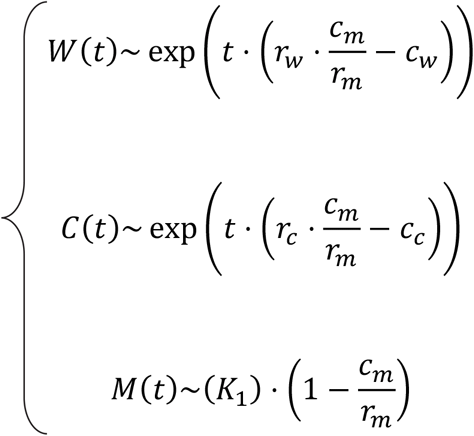

Regime R_4_ with W, M ≪ C∼ Θ(K_2_)

In this case, the equations become

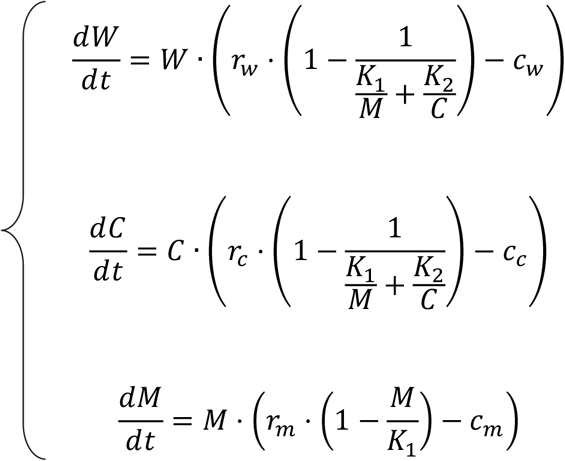

Thus, we have

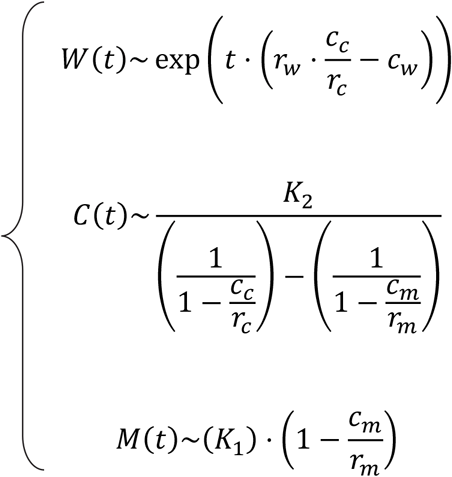

### Kinetic parameter estimation

Below we describe the different methods use to estimate the value of the different kinetic parameters of the model from the competition data (see Figure 1). Table 1 recapitulates these results. These results suggest that *S*.Tm^Comp^ has a higher replication rate than the *S*.Tm^WT^ and that vaccination induces a higher clearance rate for *S*.Tm^WT^ strain.

#### Replication rates

The replication rates can be estimated from pAM34 experiments (see Fig. S2). pAM34 has been engineered to replicate only in the presence of IPTG and pAM34 carrying bacteria can be tracked by ampicillin resistance encoded on the plasmid. Therefore, the final proportions of plasmid-carriers *y*_0_ can be linked to the number of generations, *g*_0_:

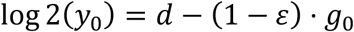

where 2^d^ and ε are related to the initial number of plasmid copies and to the residual replication rate, respectively. The measurement of the *in vitro* proportion of plasmid-carriers *y*_0_ for successive dilutions enables the estimation of *d* and ε. Table S4 shows the estimation of *d* and ε values from *in vitro* data.

**Table S4:** Estimation of *d* and ε from *in vitro* data. The value in brackets for *S*.Tm^Comp^ gives the day when *S*.Tm^Comp^ was given as compared to *S*.Tm^WT^.

| Strain | $d$ | $\varepsilon$ |
| --- | --- | --- |
| $S.Tm^{WT} Kan^R$ | 2.15 | 0.11 |
| $S.Tm^{WT} Cm^R$ | 1.9 | 0.1 |
| $S.Tm^{Comp} Cm^R$ (D0) | 3.43 | 0.22 |
| $S.Tm^{Comp} Cm^R$ (D-3) | 3.38 | 0.16 |

The replication rates can then be obtained from *in vivo* competition data. Indeed, in these experiments, each strain followed an exponential growth regime at short times. In this regime, we can estimate the number of generations *g* at time *t* given access to the maximum replication rate *r* as

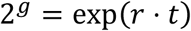

Here *g* was estimated from the *in vivo* proportion of plasmid-carriers at 12 h post infection and from the *in vitro* estimation of *d* and ε. Table S5 shows the growth rate calculated for each strain for the different competition conditions.

**Table S5:** Estimation of the growth and clearance rates from *in vivo* competition data at *t* = 12 h. For each experiment *k*, we have expressed the value of the growth rate *r*_k_ as 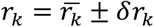 with 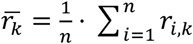 and 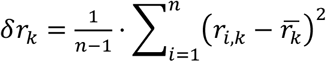. The index *n* (column #) corresponds to the number of mice having not out-diluted plasmids at *t* = 12 h. Initial *S*.Tm^WT^ Kan^R^/*S*.Tm^WT^ Cm^R^ CFU: 6000-7500. Initial *S*.Tm^WT^ Kan^R^/*S*.Tm^Comp^ Cm^R^ CFU: 3600.

| Group | Strain | Growth rate (day<sup>-1</sup>) | # mice |
| --- | --- | --- | --- |
| Naive w/o $S.Tm^{Comp}$ | $S.Tm^{WT}$ Kan<sup>R</sup> | 34.4±2.4 | 3 |
| | $S.Tm^{WT}$ Cm<sup>R</sup> | 32±3.3 | 2 |
| Naive + $S.Tm^{Comp}$ | $S.Tm^{WT}$ Kan<sup>R</sup> | 31.8±4.8 | 3 |
| | $S.Tm^{Comp}$ Cm<sup>R</sup> | 36±4.3 | 3 |
| Vacc. w/o $S.Tm^{Comp}$ | $S.Tm^{WT}$ Kan<sup>R</sup> | 30.4 | 1 |
| | $S.Tm^{WT}$ Cm<sup>R</sup> | 25.9±3.6 | 2 |
| Vacc. + $S.Tm^{Comp}$ | $S.Tm^{WT}$ Kan<sup>R</sup> | 33.7±3.7 | 3 |
| | $S.Tm^{Comp}$ Cm<sup>R</sup> | 38.7±0.4 | 3 |

#### Clearance rates

To estimate the clearance rates, we have used the fact that the size of *S*.Tm^WT^ starts to decrease exponentially after a few days. For the naive group, we can thus write

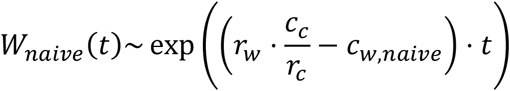

that is

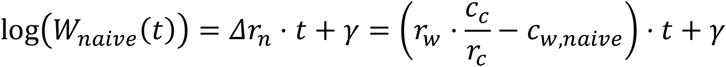

This holds approximatively from day 1 to day 8 post infection, after which the bacterial number plateaus. Here we suppose that the clearance rates of *S*.Tm^WT^ and *S*.Tm^Comp^ are similar in the naïve case, that is c_w,naïve_ = c_c_. The underlying hypothesis is that the clearance rates of *S*.Tm^WT^ and *S*.Tm^Comp^ are only different in the vaccinated group due to enchained growth of *S*.Tm^WT^. Thus, the slope 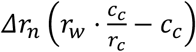 can be estimated from the *in vivo* CFUs. If we use the replication rates estimated above and make the assumption that the clearance rate of *S*.Tm^Comp^ and *S*.Tm^WT^ are the same in the naive case, we get the naïve clearance rate as

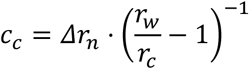

Similarly for the vaccinated group from day 1 to day 3 post infection, we can write

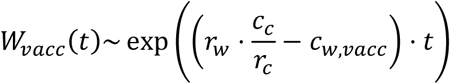

that is

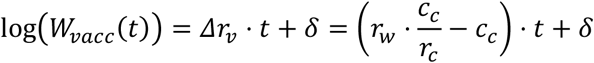

Again *Δr*_*v*_ can be estimated from the data and using the value of *c*_*c*_ obtained from the naive case, we get an estimation for the clearance rate of *S*.Tm^WT^ in the vaccinated case:

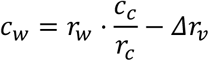

Table S6 shows the values of *c*_*w*_ and *c*_*c*_ estimated with this method.

**Table S6:** Clearance rate values.

| $r_w$ (day<sup>-1</sup>) | $r_c$ (day<sup>-1</sup>) | $c_w$ (day<sup>-1</sup>) | $c_c$ (day<sup>-1</sup>) |
| --- | --- | --- | --- |
| 33.7 | 38.7 | 6.4 | 5.9 |

#### Carrying capacities

Three carrying capacities need to be estimated. K1 can be obtained from the following expression:

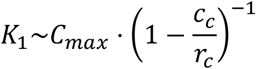

where *C*_*max*_ is the size of the *S*.Tm^Comp^ population at its maximum (typically at *t* = 24 h) in the vaccinated group.

For K_2_, we have

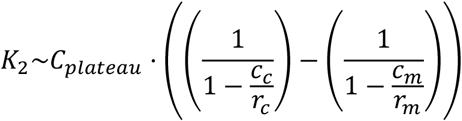

where *C*_*plateau*_ is the mean size of the *S*.Tm^Comp^ population when it reaches a plateau around 7-10 days post infection. This estimation leads to

K_1_ = 5.9·10^9^ and K_2_ = 2.3·10^4^. The value of K_3_ needs to be determined by curve fitting (see below).

#### Microbiota kinetic parameters: ratio between clearance and replication rate

The ratio between clearance and replication rate can be estimated as follows:

In the case K_3_ ≪ K_1_ when the microbiota has recovered (that is M ≫ C,W) and where C ≫ K_2_, we have

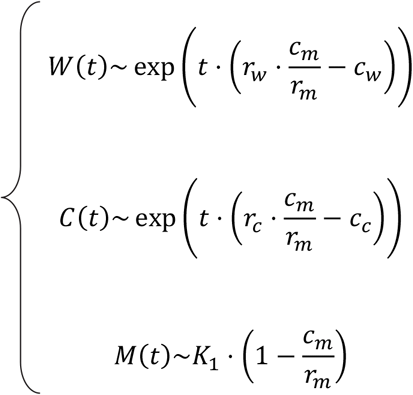

where r_m_ and c_m_ are the replication rate and the clearance rate of the microbiota, respectively.

We thus have

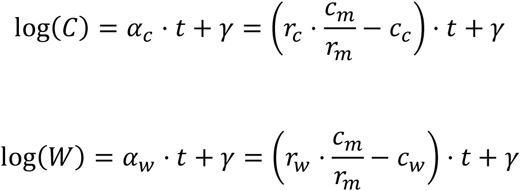

where the value of the slopes α_c_ and α_w_ can be estimated experimentally and gives access to the ratio between clearance and replication rate of the microbiota, 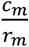. The ratio was calculated as

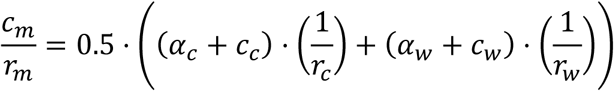

#### Data fitting

Three kinetic parameter values still need to be estimated: the size of the microbiota at *S*.Tm infection, m_0_, the microbiota replication, r_m_ (its clearance rate is then directly obtained from the ratio estimated above) and the value of the carrying capacity K_3_.

These parameters were estimated by fitting the model to the *S*.Tm^WT^ and *S*.Tm^Comp^ data (vaccinated group) with a least squares method (see Fig. 2A for the best fit and Table S7 for the corresponding values).

**Table S7:** Parameter values obtained from data fitting. The fit was performed with the nonlinear least-squares solver lsqcurvefit of MATLAB R2015a. corresponds to the value of the squared 2-norm of the residual ∑*(f(ρ, xdata) -* log_10_(*ydata*))^2^ with p = r_m_ and where *f* is the log_10_ of the solution of the ODE system. The algorithm was run for m_0_ values ranging between 10^5^ and 10^7^. The algorithm was run for K_3_ values ranging between 10^7^ and 5.10^8^.

| $m_0$ (CFUs g<sup>-1</sup> feces) | $r_m$ (day<sup>-1</sup>) | $K_3$ | Res. norm |
| --- | --- | --- | --- |
| $10^5$ | 18.7 | $10^7$ | 4 |

### Extinction probability and extinction time

It is also important to evaluate the time needed for the extinction of the *S*.Tm^WT^ population. For the sake of simplicity, we only consider cases where the initial number of newly introduced bacteria is large, that is W_0_ > 100. In this case, the dynamics of the invasion process can be divided into two time windows (Fig. 2). The first time window starts with the inoculation and ends when the number of *S*.Tm^WT^ bacteria becomes < 100: all populations are thus large so that time lapse spent in this time window is evaluated by integrating the deterministic equations

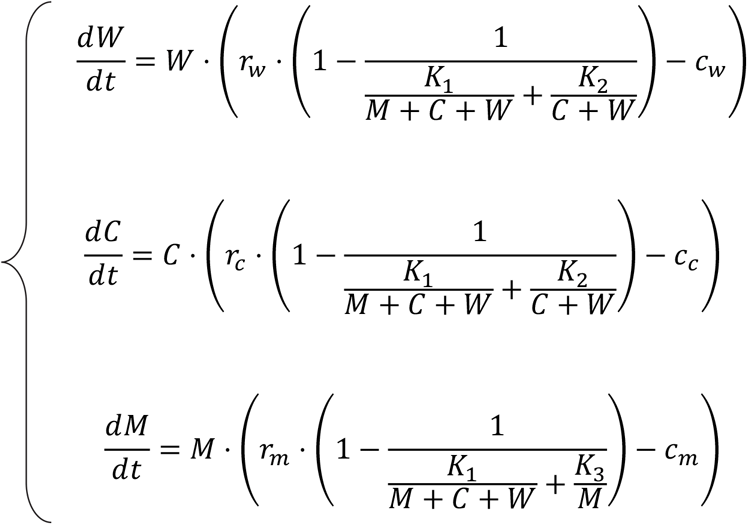

until the time τ when W (τ) = 100. The second time window starts when the resident population becomes < 100 and ends with its extinction: a stochastic description is used here because of the small size of the resident population. The distribution of the times needed to go from W = 100 to extinction can be obtained numerically with the Gillespie algorithm.

## Supplementary Figures

**Figure S1.**
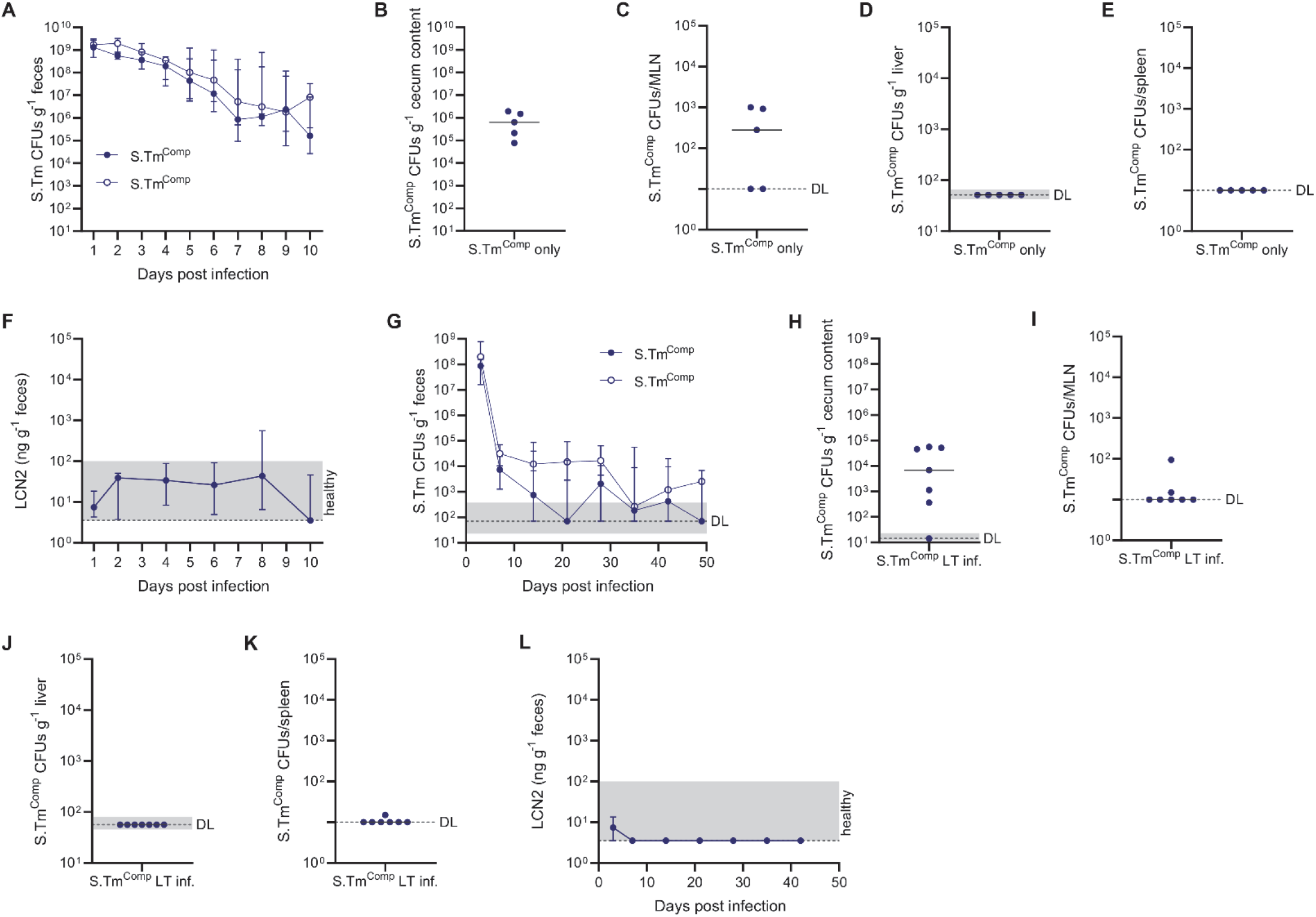
*S*.Tm^Comp^ is not pathogenic in 129S6/SvEv and C57BL/6J mice. (**A-F**) 129S6/SvEv mice were pretreated with streptomycin and infected with a total of 10^5^ of a 1:1 mixture of two isogenic *S*.Tm^Comp^ strains and colonization was followed for 10 days. *S*.Tm^Comp^ CFUs were determined by selective plating in feces (**A**) cecum content (**B**), MLN (**C**), liver (**D**) and spleen (**E**). (**F**) Intestinal inflammation was determined by measuring fecal lipocalin-2. (**G-L**) C57BL/6 mice were pretreated with streptomycin and infected with a total of 10^4^ of a 1:1 mixture of two isogenic *S*.Tm^Comp^ strains and colonization was followed for 7 weeks. *S*.Tm^Comp^ CFUs were determined by selective plating in feces (**G**) cecum content (**H**), MLN (**I**), liver (**J**) and spleen (**K**). (**L**) Intestinal inflammation was determined by measuring fecal lipocalin-2. N = 5-7 mice. Solid lines depict the median, error bars the interquartile range. Dotted lines show the detection limit and the shaded area the range for cases in which the detection limit is dependent on sample weight. Statistics were performed by an unpaired two-tailed t-test on log-normalized area under the curve (AUC) (A, G). CFU, colony forming unit; LCN2, lipocalin-2; MLN, mesenteric lymph node.

**Figure S2.**
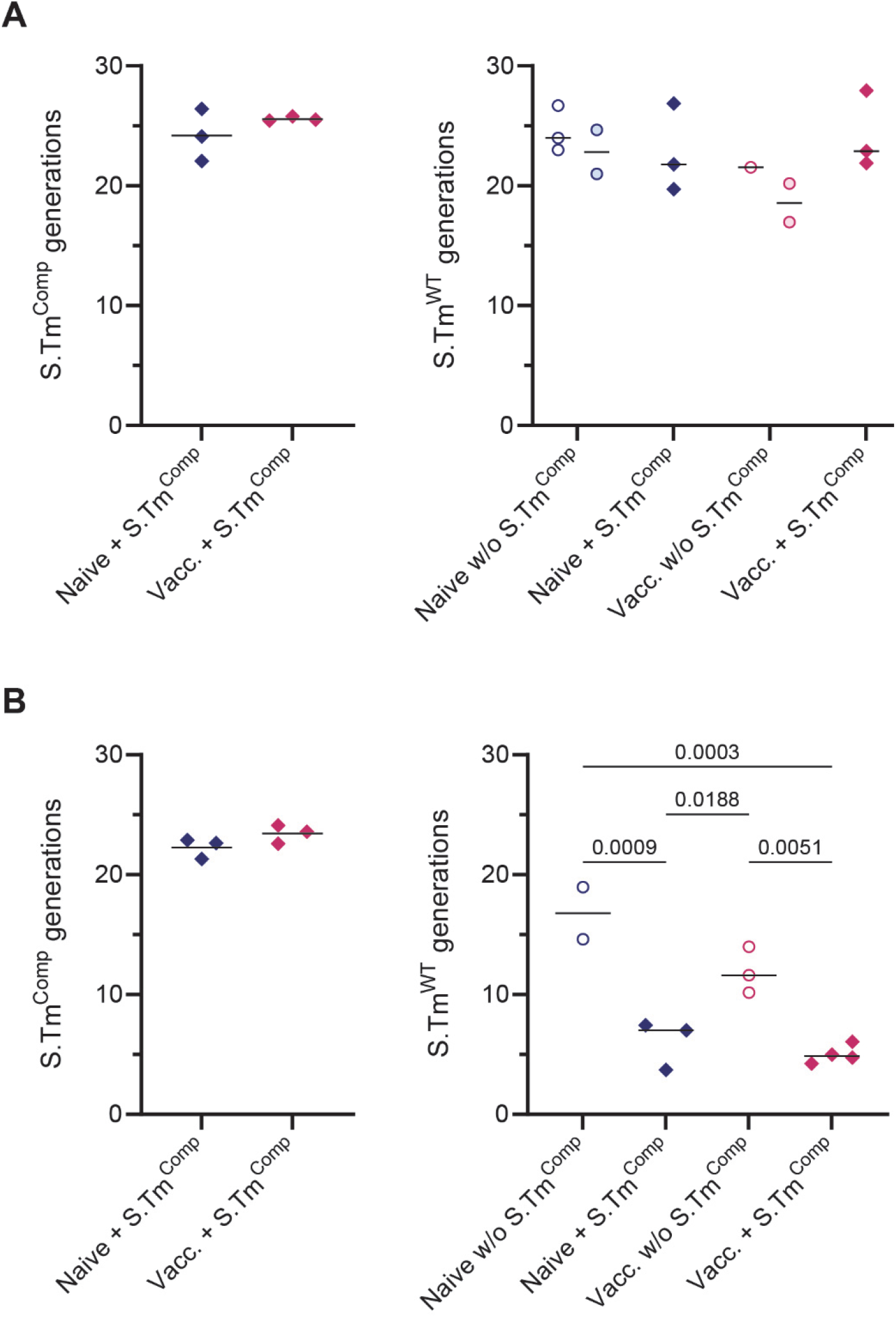
Bacterial generations *in vivo* at 12 hours post infection. (**A**) PBS (blue symbols) or PA-*S*.Tm-vaccinated (pink symbols) 129S6/SvEv mice were pretreated with streptomycin and infected with a total of 10^4^ of a 1:1 mixture of two isogenic *S*.Tm^WT^ strains (circles) or *S*.Tm^WT^ and *S*.Tm^Comp^ (*S*.Tm^*hilD ssaV oafA*^, filled diamonds). Number of generations 12 hours post infection was estimated based on the loss of pAM34. Open and filled circles show the two isogenic *S*.Tm^WT^ strains. (**B**) PBS (blue symbols) or PA-*S*.Tm-vaccinated (pink symbols) 129S6/SvEv mice were pretreated with streptomycin and infected with 1·10^6^ *S*.Tm^WT^. Two groups were pre-colonized with 5·10^3^ *S*.Tm^Comp^ 3 days before infection. Number of generations 12 hours post infection was calculated based on the loss of pAM34. Solid lines depict the mean. Statistics were performed by one-way ANOVA.

**Figure S3.**
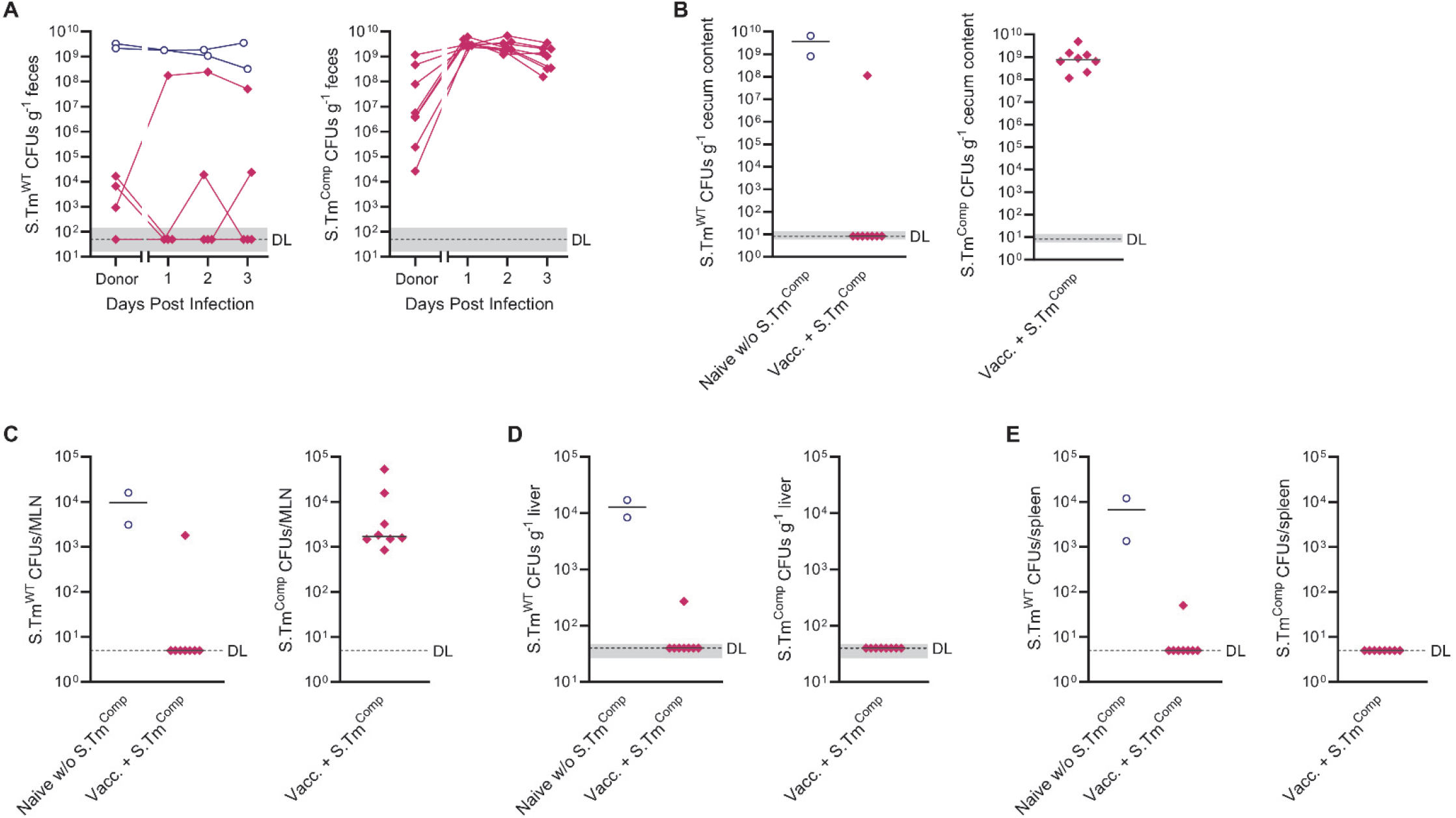
Vaccination together with *S*.Tm^Comp^ a priori colonization prevents *S*.Tm^WT^ transmission. Naïve 129S6/SvEv mice were pretreated with streptomycin and a FMT was performed with feces collected from 9 days post infection of untreated (blue circles) or vaccinated and *S*.Tm^Comp^ pre-colonized mice (pink diamonds; see Fig. 3). *S*.Tm^WT^ and *S*.Tm^Comp^ CFUs were determined by selective plating in feces (**A**) cecum content (**B**), MLN (**C**), liver (**D**) and spleen (**E**). Pooled data from two independent experiments with switched antibiotic resistances (n = 2 or 8 mice/group). Solid black lines depict the median. Dotted lines show the detection limit and the shaded area the range for cases in which the detection limit is dependent on sample weight. CFU, colony forming unit; FMT, fecal microbial transplant; MLN, mesenteric lymph node.

**Figure S4.**
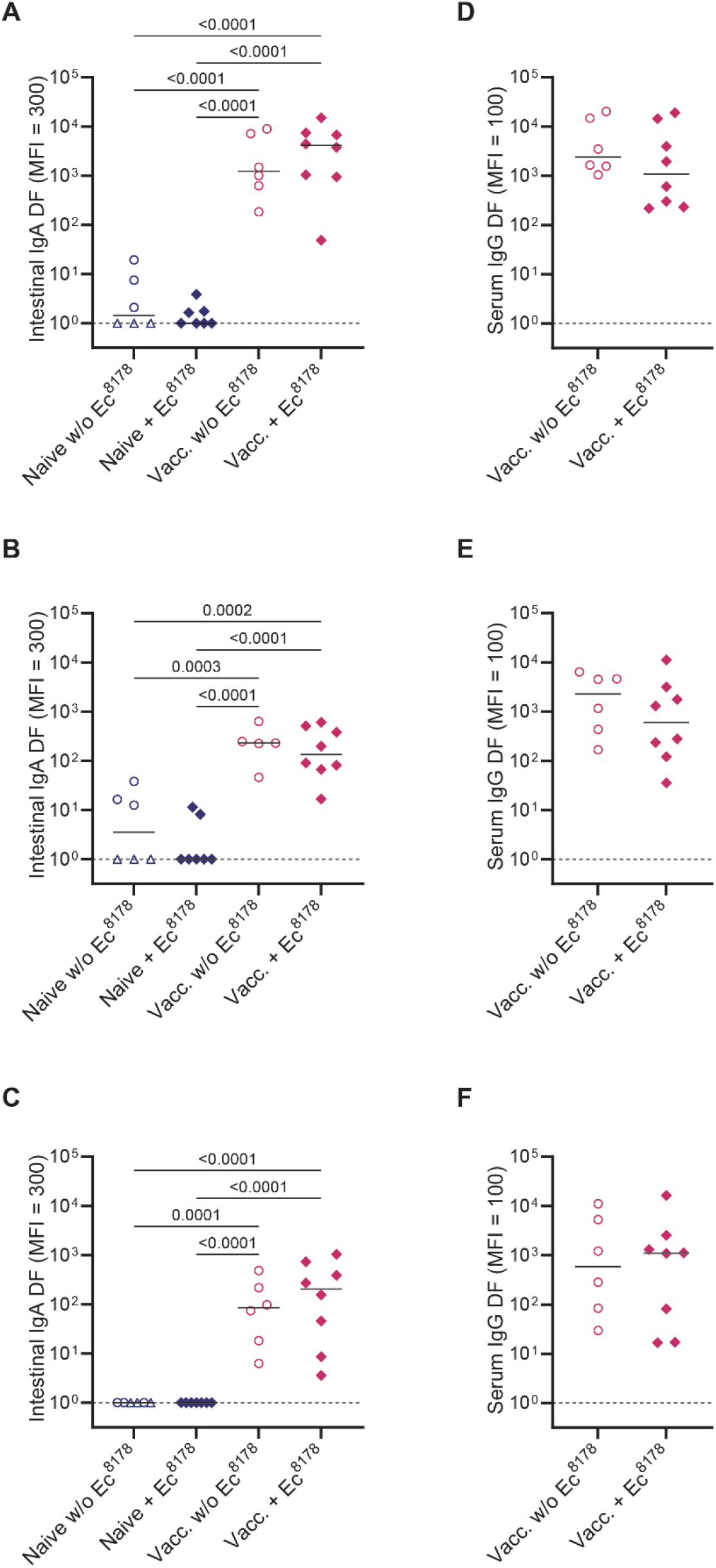
Vaccination with the EvoTrap vaccine generates an intestinal IgA and serum IgG response against all *S*.Tm O-antigen variants. 129S6/SvEv mice were mock-vaccinated with PBS (blue symbols) or EvoTrap-vaccinated (pink symbols) and later infected with 1.10^6^ *S*.Tm^WT^ with or without pre-colonization with 5.10^3^ *E. coli* 8178. 10 days after *S*.Tm^WT^ infection *S*.Tm specific antibody responses were determined in intestinal lavage (**A-C**) and serum (**D-E**) by flow cytometry. (**A+D**) *S*.Tm O:4,12-0. (**B+E**) *S*.Tm O:4[5],12-2. (**C+F**) *S*.Tm O:4,12-2. Pooled data from two independent experiments (n = 6-8 mice/group). Solid lines depict the. Dotted lines show the detection limit. Open triangles show mice that had to be euthanized prematurely due to excessive weight loss (≥ 15%) or disease symptoms. Statistics were performed one-way ANOVA on log-normalized data (A-C). Where only two groups were compared, an unpaired two-tailed t-test on log-normalized data was done (D-F). DF, dilution factor; MFI, median fluorescence intensity.

**Figure S5.**
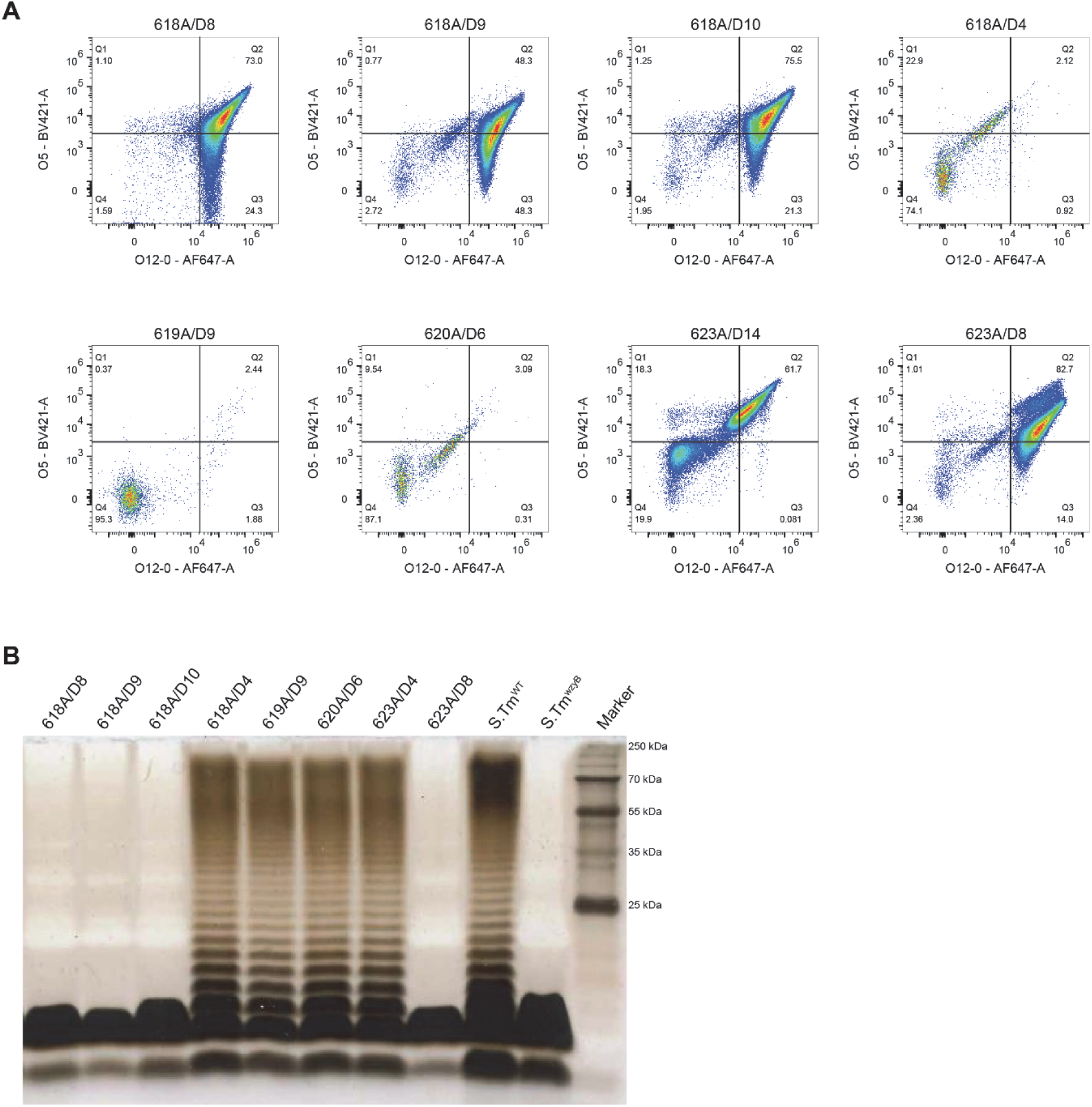
Vaccination with the EvoTrap vaccine leads to emergence of *S*.Tm^WT^ clones with short O-antigen. 129S6/SvEv mice were EvoTrap-vaccinated and later infected with 1.10^6^ *S*.Tm^WT^ with or without pre-colonization with 5.10^3^ *E. coli* 8178. *S*.Tm^WT^ clones were isolated from feces from 4 days post infection onwards. (**A**) *S*.Tm^WT^ clones were monitored for decreased O:5 and O:12-0 staining intesity by flow cytometry. (**B**) Silver-stained gel of LPS from selected *S*.Tm^WT^ clones from 4 different EvoTrap vaccinated mice.

**Figure S6.**
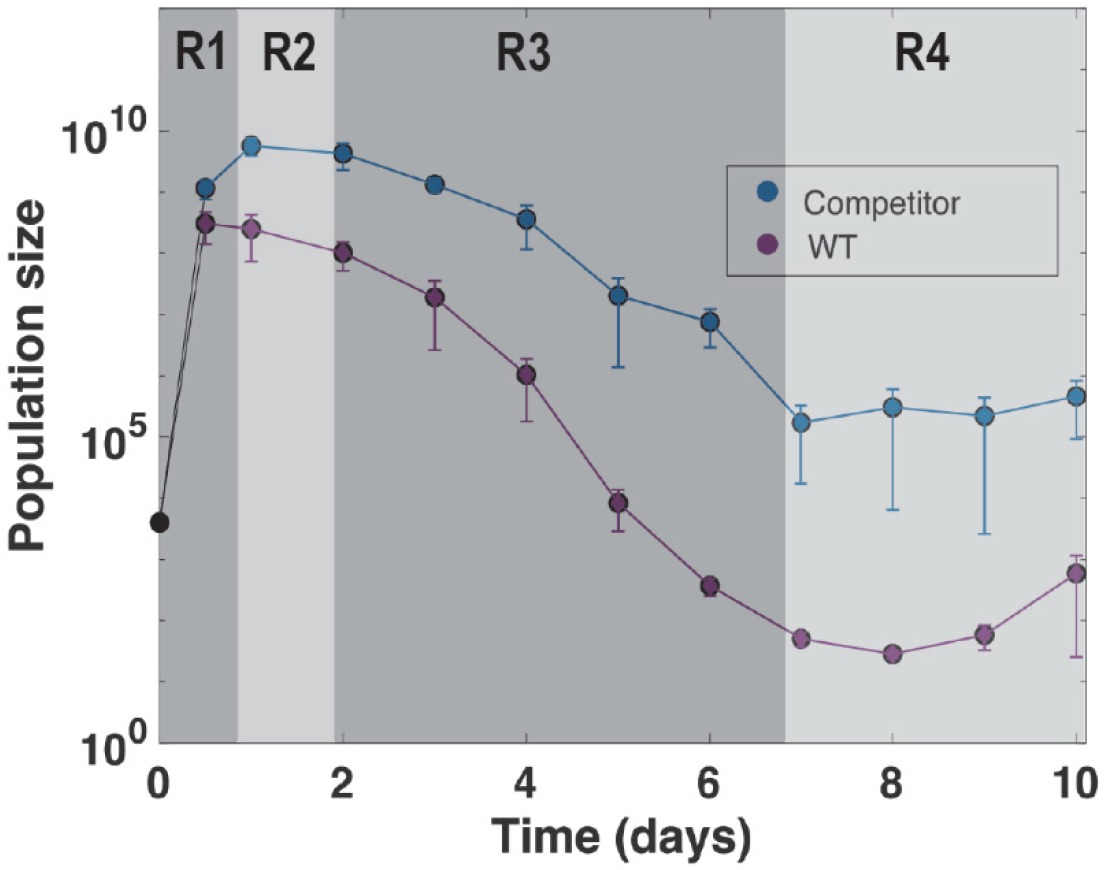
Different regimes used for mathematical modelling. PA-*S*.Tm-vaccinated 129S6/SvEv mice were pretreated with streptomycin and infected with a total of 10^4^ of a 1:1 mixture of *S*.Tm^WT^ (blue symbols) and *S*.Tm^Comp^ (purple symbols). R_1_ defines the time window where *S*.Tm^WT^, *S*.Tm^Comp^ and the microbiota are all below their carrying capacity. R_2_ is defined by *S*.Tm^Comp^ being more abundant than *S*.Tm^WT^ and the microbiota, and being in the order of magnitude of K_1_. R_3_ is defined by the microbiota being more abundant than *S*.Tm^WT^ and *S*.Tm^Comp^ but *S*.Tm^Comp^ is still present in higher numbers than its carrying capacity in equilibrium K_2_. R_4_ is defined by the microbiota being much more abundant than *S*.Tm^WT^ and *S*.Tm^Comp^ and *S*.Tm^Comp^ being in the order of magnitude of K_2_.

